# Thiol Scarcity in Cerebrospinal Fluid Renders Leptomeningeal Acute Lymphoblastic Leukaemia Therapeutically Vulnerable to Ferroptosis

**DOI:** 10.64898/2025.12.15.693383

**Authors:** Nikolai Gajic, Rhona Christie, Shirley Lam, Tobias Ackermann, Ekaterini Himonas, Jonathan Josephs-Spaulding, Anand Manoharan, Victoria Assmann, Jordan Martin, Alexander Collings, Karen Dunn, Engy Shokry, Alejandro Huerta Uribe, Sharon Burns, David O’Connor, Marc R. Mansour, David Sumpton, G. Vignir Helgason, Johan Vande Voorde, Saverio Tardito, Stephen WG Tait, Christina Halsey

## Abstract

The leptomeninges present a challenging tumour microenvironment with cells receiving low levels of nutrients and oxygen from cerebrospinal fluid (CSF), however these metabolic constraints are yet to be exploited therapeutically. Central nervous system (CNS) relapse in acute lymphoblastic leukaemia (ALL) remains a formidable clinical challenge because the leptomeningeal niche restricts drug penetration and immune surveillance. Current CNS-directed treatments rely on neurotoxic intrathecal chemotherapy, underscoring the urgent need for novel targeted strategies. Here, we uncover a profound niche-specific metabolic vulnerability in CNS-resident ALL cells, characterised by an obligate reliance on LRP8-mediated selenium uptake to sustain selenocysteine biosynthesis and GPX4 activity under profound glutathione limitation. We show that scarcity of thiols and cystine in CSF creates an inherently pro-ferroptotic microenvironment. Interference with selenocysteine biosynthesis under these conditions induces synthetic lethality in both *in vitro* and *in vivo* CNS-ALL models. This vulnerability is exploitable both by genetic targeting of the selenocysteine biosynthesis pathway and, notably, through repurposing the FDA-approved agent Auranofin, which disrupts selenium utilisation, induces lipid peroxidation, and demonstrates CNS-specific anti-leukaemic efficacy with excellent tolerability *in vivo*. These findings identify a novel mechanistically grounded approach, leveraging features of the unique leptomeningeal microenvironment to selectively kill invading cells, with potential implications for all leptomeningeal-tropic malignancies.

**Graphical Abstract:**
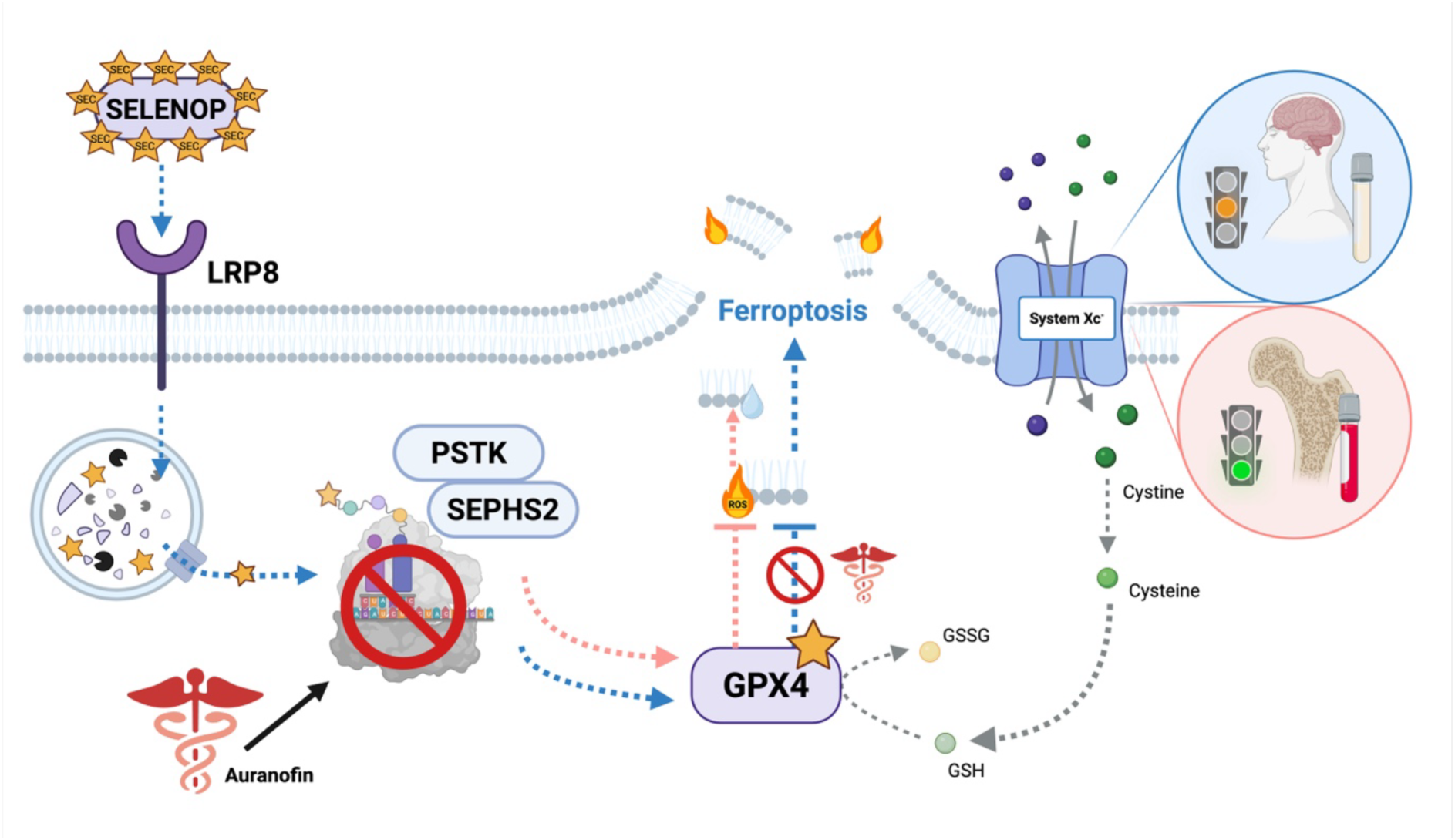
The leptomeningeal tumour microenvironment is intrinsically pro-ferroptotic due to low concentrations of free thiols and cystine in cerebrospinal fluid. ALL cells in this niche exhibit disrupted glutathione metabolism and a novel dependency on LRP8-mediated SELENOP uptake to sustain GPX4 levels and thus protect themselves from spontaneous ferroptotic cell death. Perturbation of selenocysteine biosynthesis via genetic inhibition SEPHS2 or LRP8 leads to synthetic lethality in leukaemic cells exposed to CSF *in vitro* and *in vivo* – an effect phenocopied by use of Auranofin.

## Introduction

Cerebrospinal fluid (CSF) is an ultrafiltrate of plasma that plays important roles in mechanical protection of the brain and spinal cord^1^ as well as supporting central nervous system (CNS) immune surveillance and removal of metabolic waste to protect brain function^2^. This highly specialised fluid has a distinct composition with low levels of lipids, proteins and energy sources compared to plasma^3^. CNS leukaemia and other leptomeningeal metastasis reside within the leptomeninges bathed in CSF. Among human malignancies, childhood acute lymphoblastic leukaemia (ALL) is unique in that every patient receives therapy specifically targeting leptomeningeal-resident cells^4^. CNS-directed therapies, such as methotrexate (MTX), are associated with substantial neurotoxicity, including seizures and persistent cognitive impairments^5^. Moreover, there are no novel agents for CNS relapse and immunotherapies are less effective in this compartment^6^. This underscores the urgent need to deepen our mechanistic understanding of leukaemic cell adaptation within the CNS niche to identify new therapeutic targets. We hypothesised that exposure to CSF would drive unique metabolic adaptations in highly proliferative cancer cells with significant nutritional demands. Consistent with this, we previously identified Stearoyl-CoA desaturase (SCD) alongside a distinct fatty acid and cholesterol synthesis signature^7^ in CNS-ALL, suggesting metabolic vulnerabilities as potential therapeutic targets. A similar signature has also been observed in other leptomeningeal cancers^8^ suggesting a universal metabolic bottleneck for proliferating cells in this niche.

One potential reason for SCD dependence in the CNS could be its role in protecting cells from ferroptosis. Ferroptosis is a non-apoptotic iron dependent form of cell death characterised by overwhelming lipid peroxidation. Ferroptosis is kept in check by the selenoprotein glutathione peroxidase 4 (GPX4), which uniquely detoxifies phospholipid peroxides by reducing them to non-reactive lipid alcohols. This protective activity depends critically on the presence of the rare amino acid selenocysteine at the active site of GPX4, and the availability of glutathione (GSH) as a reducing cofactor. Further protection from ferroptosis is afforded by SCD driven production of non-oxidisable monounsaturated fatty acids (MUFAs), to prevent the characteristic chain reaction of lipid peroxidation across cell membranes that leads to cell death. Targeting ferroptosis shows promise for cancer treatment in specific contexts^9^^10^, however, additional investigation is required to fully clarify its therapeutic potential^11^^12^.

Here, we uncover a profound metabolic vulnerability to ferroptosis driven by CSF exposure in the leptomeningeal niche. We show that low cystine and free thiol levels in CSF disrupt GSH synthesis and prime leukaemic cells for lipid peroxidation and ferroptotic cell death. We identify significant alterations in the expression of key selenocysteine biosynthesis genes and ferroptosis regulators in primary patient-derived CNS-ALL blasts. Within the CNS, ALL cells are dependent on scavenging of scarce selenium reserves to generate enough functional GPX4 to prevent ferroptosis. We demonstrate that targeting selenium uptake, through multiple means including repurposing of the FDA approved drug Auranofin, significantly reduces CNS leukaemia burden in ALL xenograft cell line and patient-derived models *in vivo*. These findings establish ferroptosis as a previously unrecognised, leptomeningeal niche-dependent therapeutic vulnerability, which may be leveraged through drug repurposing.

## Results

### Ferroptosis-Related Gene Expression is Altered in CNS-Resident ALL Cells

We have previously demonstrated a selective reliance on SCD in acute lymphoblastic leukaemia (ALL) blast cells residing in the central nervous system (CNS)^7^. Given SCD’s now well-established role as a ferroptosis regulator^13^^14^, we asked whether cell death observed previously was inflicted via caspases or through lipid peroxidation. Interestingly, we observed that reduced viability associated with SCD inhibition could be wholly rescued by Ferrostatin-1, a specific inhibitor of membrane lipid peroxides, with no significant rescue with QVD, a pan-caspase inhibitor (**Figure 1A**). These data indicate that SCD inhibition can induce ferroptosis in ALL cells in a context dependent manner. We next explored existing patient datasets to establish whether, alongside *SCD*, the expression of other ferroptosis related genes were altered (**Figure 1B**). To this end, we compared the transcriptomic profiles of patient-derived CNS-ALL with those of bone marrow ALL^15^. This revealed transcriptional changes of genes involved in the membrane incorporation of MUFAs (**Figure 1C**), selenium uptake and utilisation (**Figure 1D**), Ferroptosis Suppressor Protein 1 (*FSP1*) (**Figure 1E**), and iron uptake (**Figure 1F**) consistently suggesting a ferroptosis-protective metabolic reprogramming in the CNS resident cells. Given the robust and consistent regulation of selenium-handling genes (**Figure 1D**) we investigated the availability of selenium to ALL cells in CSF. A literature review on selenium speciation in mammalian CSF versus plasma revealed that selenium levels are markedly lower in CSF, with Selenoprotein P (SELENOP) identified as the main transporter of bioavailable selenium in the CNS (**Figure 1G, Supplementary Table 1**). Optimal selenium utilisation depends on the availability of thiol-containing proteins, particularly glutathione (GSH)^16^. To assess thiol availability in CSF, we employed Ellman’s assay, an established test for thiol groups^17^. Thiol levels were found to be markedly lower in patient CSF compared with their paired plasma, in agreement with previously published literature^18^ (**Figure 1H**). GSH is a tripeptide composed of the amino acids cysteine, glycine and glutamate, with cystine serving as the most critical precursor, since its uptake and reduction to cysteine often represents the rate-limiting step for GSH synthesis^19^. To assess whether this constraint is further exacerbated in ALL, we compared CSF cystine levels in healthy individuals with those measured in children with ALL. Strikingly, cystine was nearly undetectable in ALL patient CSF, indicating aggressive depletion of the already limited cystine pool (**Figure 1I, Supplementary Table 2**)^20^. Collectively, these data underscore the severe nutrient constraints faced by ALL cells within the CNS and highlight cystine and selenium scavenging as potential metabolic adaptations that support their survival in this nutrient-poor microenvironment.

**Figure 1:**
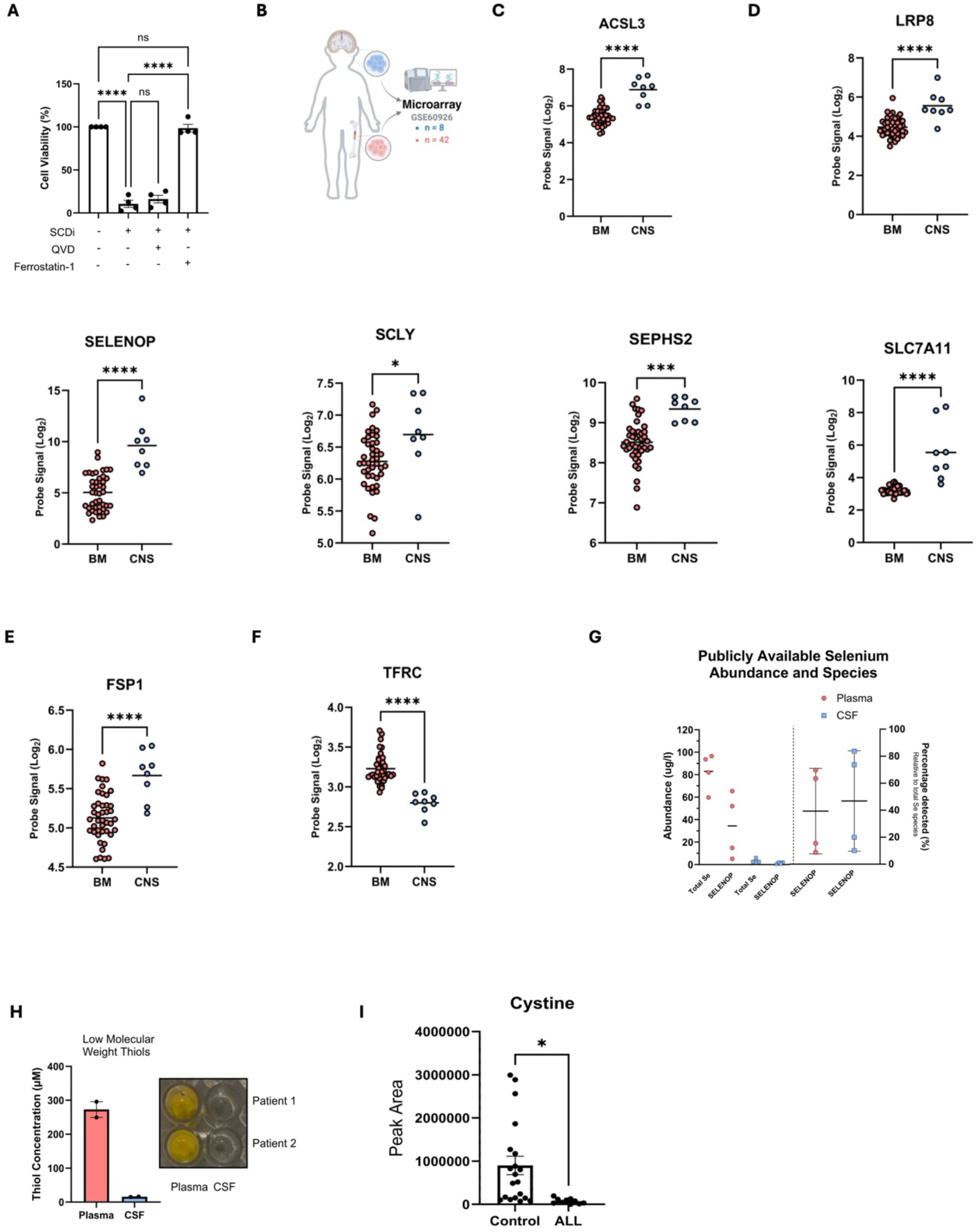
Ferroptosis-Related Gene Expression is Altered in CNS-Resident ALL Cells. (**A**) SEM cells treated with 2.5μM SW203668 for 72 hours in delipidated serum 10% / 90% RPMI with or without 2μM Ferrostatin-1 or 10μM QVD as indicated. N=4 independent experiments. P-values refer to a one-way ANOVA test with Tukey’s multiple comparisons test. Error bars represent mean ± SEM. **(B)** Schematic depicting publicly available dataset GSE60926. Created with BioRender.com. **(C-F)** Probe signals or normalised counts from dataset GSE60926 comparing **(C)** ACSL3, **(D)** LRP8, SELENOP, SCLY, SEPHS2, and SLC7A11 (System Xc^-^), **(E)** FSP1, **(F)** TFRC gene expression in CNS (blue circle, n = 8) vs BM (red circle, n = 42) derived leukaemic cells. P-values calculated using an unpaired two-tailed Student’s t-test. **(G)** Left panel: Summary of selenium and SELENOP levels in plasma and cerebrospinal fluid (CSF) according to published values. Right panel: Proportion of SELENOP relative to total selenium in plasma versus CSF. A reference list and individual means of detection are provided in **Supplementary Table 1**. **(H)** Left panel: Quantification of low molecular weight thiols in patient samples using Ellman’s reagent. Right panel: Representative images of wells with yellow-coloured product formed after mixing plasma or CSF with Ellman’s reagent. Error bars represent mean ± SEM. **(I)** LC-MS peak area values of cystine in CSF of healthy control subjects (n = 20) or patients with Acute Lymphoblastic Leukaemia (n = 10). P-values calculated using unpaired two-tailed Student’s t-test. Error bars represent mean ± SEM. * ≤ 0.05 *** ≤ 0.001 **** ≤ 0.0001

### CNS-Resident ALL Cells Display Disrupted Glutathione Metabolism

Lipid peroxidation is a hallmark of ferroptosis^21^. Given the profound changes in gene expression and ferroptosis-associated metabolite abundance in the CNS niche, we next used the C11-BODIPY sensor^22^ to investigate whether there were differences in basal levels of lipid peroxidation between ALL cells resident in the bone marrow or CNS compartment (**Figure 2A**). Lipid peroxidation levels were significantly higher in CNS-resident ALL cells across multiple models, including cell lines and both B– and T-ALL patient-derived xenografts (PDXs) (**Figure 2B-D, Sup. 1A, B**). ChaC glutathione-specific γ-glutamylcyclotransferase 1 (CHAC1) is an ER stress inducible γ-glutamyl cyclotransferase that catalyses GSH degradation downstream of the ATF4 response^23^. CHAC1 has been previously identified as a biomarker of ferroptosis induction^24^. In line with previous reports^24^, we observed that *CHAC1* expression was selectively upregulated by cystine deprivation and by the System Xc⁻ inhibitor Erastin, but not by other stressors such as dual BCL-2/BCL-XL inhibition, direct GPX4 inhibition, or serum/selenium withdrawal (**Figure Sup. 1C**). Using *CHAC1*, we surveyed RNA-seq datasets from cancer cells in the leptomeningeal niche, including a BCP-ALL PDX model (GSE271998; **Figure 2E**), ALL patient CNS relapse samples (GSE60296; **Figure Sup. 1D**), BCP-ALL REH cells (**Figure Sup. 1E**), a T-ALL xenograft model (**Figure Sup. 1F**), and metastatic CNS models of lung, breast, and liver cancer (**Figure Sup. 1G**). In all contexts, *CHAC1* expression was upregulated. Notably, *CHAC1* and the canonical ATF4 stress response were specifically enriched in CNS samples, but not in bone marrow or spleen from BCP ALL PDX samples (**Figure 2E, F**). Consistent with this, direct measurements revealed significantly reduced intracellular GSH in CNS-versus bone marrow–resident ALL cells (**Figure 2G).**

**Figure 2:**
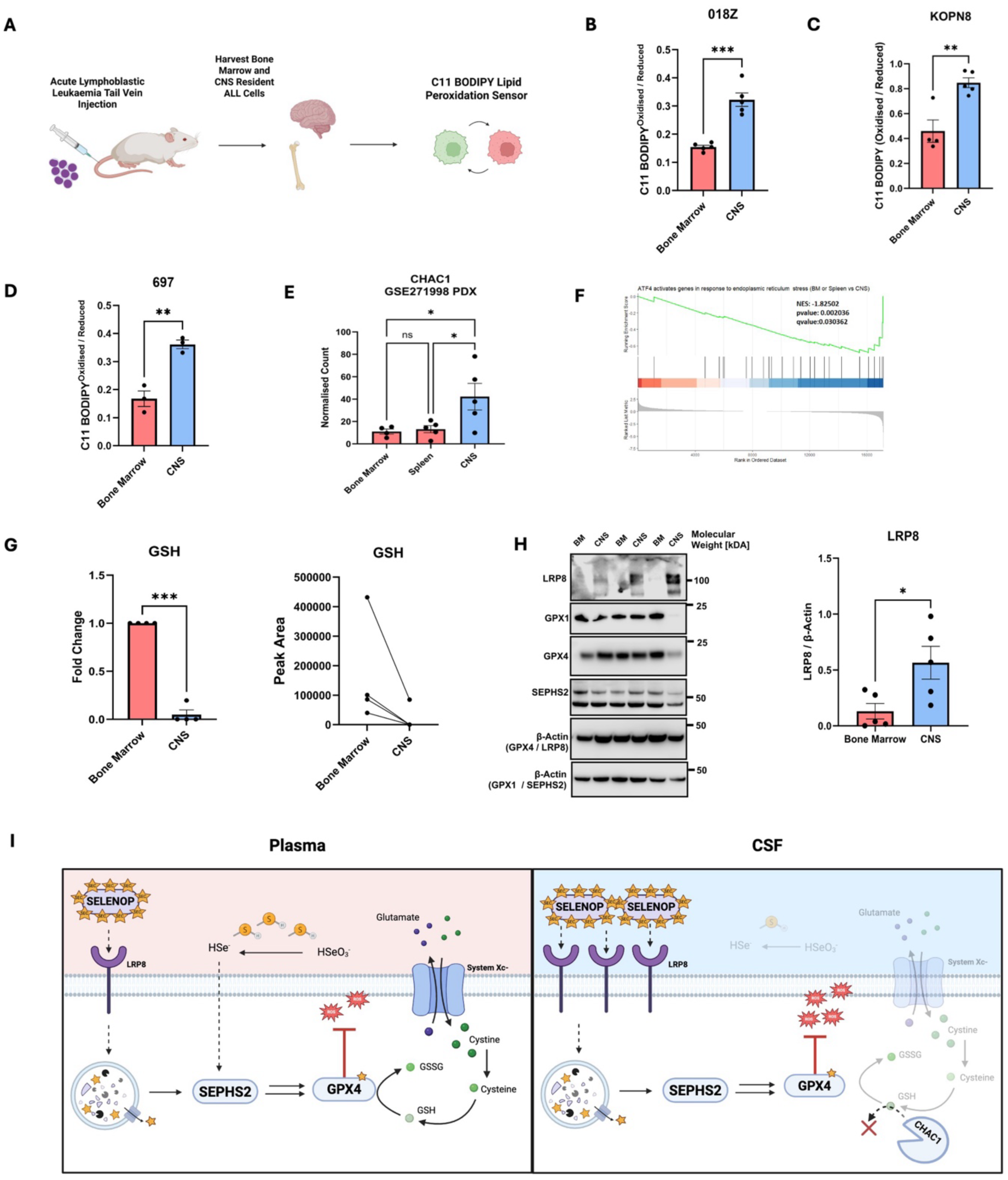
CNS-Resident ALL Cells Display Disrupted Glutathione Metabolism. (**A**) Schematic of the xenotransplantation of ALL cells into NSG mice and analysis of lipid peroxidation. Created with BioRender.com. **(B-D)** Ratio of oxidised (510 nm) to reduced (591 nm) BODIPY 581/591 C11 (lipid peroxidation sensor) in BM and CNS from **(B)** 018Z, **(C)** KOPN8 and **(D)** 697 cells retrieved from NSG mice (n = 3–5 mice, as indicated by points). P-values calculated using unpaired two-tailed Student’s t-test. Error bars represent mean ± SEM. **(E)** Gene expression of CHAC1 from GSE271998 in patient derived TCF3::PBX1+ BCP-ALL transplanted into NSG mice (n = 5) and isolated from BM, spleen, and CNS. P-values refer to a one-way ANOVA test with Tukey’s multiple comparisons test. Error bars represent mean ± SEM. **(F)** Enrichment plot of REACTOME term: ATF4 Stress Response, from RNA-seq data (GSE271998) in patient derived TCF3::PBX1+ BCP-ALL transplanted into NSG mice (n = 5) and isolated from BM, spleen, and CNS. **(G)** Intracellular abundance of GSH in SEM cells obtained from the BM and CNS of xenografted mice. Data represent n=4 paired BM and CNS mouse samples. P-value refers to a paired one-tailed t-test. Results are reported as relative fold change (Left panel) Peak area normalised to cell count and shown as arbitrary units per 10⁶ cells (Right panel). Error bars represent mean ± SEM. **(H)** Left Panel: Immunoblot image of LRP8 (GPX4 membrane), SEPHS2 (GPX1 membrane), GPX1, GPX4 and β-Actin protein in 859I PDX BCP-ALL cells retrieved from the CNS and BM of xenografted mice at clinical endpoint. 3/5 mice. Right Panel: Quantification of LRP8 from n = 5 mice. P-values calculated using unpaired two-tailed Student’s t-test. Error bars represent mean ± SEM. **(I)** Schematic representation of the proposed model of selenocysteine metabolism in the BM and CNS. In the plasma, abundant cystine and free thiols support GPX4 expression and activity, resulting in lower expression of LRP8 *(Left panel).* In contrast, CSF contains low levels of thiols and cystine, which leads to upregulation of LRP8 and CHAC1. This promotes increased uptake of SELENOP to counteract elevated lipid peroxidation *(Right panel)*. Created with BioRender.com. Error bars represent mean ± SEM. * ≤ 0.05 ** ≤ 0.01 *** ≤ 0.001

Selenium can be acquired through two routes, an organic pathway mediated by LRP8 and SELENOP, or an inorganic pathway dependent on GSH^25^. Given the significant reduction in intracellular GSH, and induction of its canonically partnering stress response, we asked whether this impacted intracellular selenoprotein abundance in the CNS. Despite the reported reduction in extracellular selenium availability (**Figure 1G**), the expression of GPX1 and GPX4 – proteins we found to be highly sensitive to selenium deprivation in vitro (**Figure Sup. 1H**) – remained comparable between CNS– and bone marrow-resident ALL cells (**Figure 2H**). Strikingly, LRP8 protein levels were significantly elevated in CNS-resident ALL cells (**Figure 2H, Sup 1I**), consistent with patient-derived mRNA data (**Figure 1D**). In solid tumours we detected an upregulation of Selenium Binding Protein 1 (*SELENBP1*), suggesting an alternative or complementary mechanism for adaptive selenium acquisition (**Figure Sup 1J**).

Thiol-containing molecules such as GSH and cysteine can be synthesised either from cystine via System Xc⁻ uptake, or de novo from methionine through the transsulfuration pathway^26^. To examine the relationship between LRP8 expression and thiol availability, we restricted cystine or methionine from ALL cells and monitored LRP8 protein levels. Cystine deprivation – but not methionine withdrawal – induced LRP8 expression in a dose-dependent manner (**Figure Sup. 1K, L**), indicating a compensatory shift between selenium uptake pathways under thiol limitation. These findings indicate that LRP8-mediated selenium uptake could be selectively induced by thiol depletion in the CNS. Together, these findings suggest that the CNS imposes a thiol-restricted environment on ALL cells, limiting GSH biosynthesis and driving elevated lipid peroxidation. In response, ALL cells upregulate LRP8 to enhance SELENOP uptake and sustain GPX expression, thereby protecting against further lipid peroxidation and ferroptosis (**Figure 2I**).

### Cerebrospinal Fluid Primes ALL Cells for Ferroptosis

To determine whether the lipid stress observed *in vivo* was primarily driven by CSF, ALL cells were cultured for 24 hours in patient-derived CSF, CSF supplemented with α-tocopherol (a lipophilic antioxidant) or Plasmax™ as a control. Plasmax™ represents a physiologically relevant cell culture medium designed to closely mimic the nutritional and metabolic profile of human plasma^27^. CSF exposure led to a clear induction of lipid peroxidation which could be quenched with α-tocopherol (**Figure 3A**). Supporting this observation, publicly available transcriptomic data^28^ from MOLT-4 (T-ALL) and REH (BCP ALL) cells cultured in either standard RPMI with 10% FBS or human CSF revealed a significant upregulation of *CHAC1*, indicating that CSF itself activates stress response pathways associated with GSH depletion and induces lipid peroxidation (**Figure 3B**, **Sup. 2A**). Due to limited availability of patient CSF, we utilised a custom physiological medium, termed “CSFmax,” (manuscript in preparation) to model human CSF. This medium included 2.9 nM selenite (reflecting a 10-fold reduction compared to Plasmax™), 1.25% FBS, and 3.75% delipidated FBS as the primary selenium sources. As observed with patient CSF, CSFmax induced substantial lipid peroxidation, which could be effectively quenched through α-tocopherol supplementation (**Figure 3C**, **Sup. 2B**) and induced *CHAC1* mRNA **(Figure 3D, Sup. 2C)**. Next, we tested the ferroptosis sensitivity to the ferroptosis inducer 1S-3R, RSL3 (GPX4 inhibitor, RSL3) (**Figure 3E**) of 697 (a B-cell precursor cell line)^29^ and CCRF-CEM (a T-ALL line)^30^ cultured in CSFmax. The treatment with RSL3 (GPX4i) revealed a marked increase in ferroptotic sensitivity in CSFmax compared to Plasmax™, an effect that was rescued by the ferroptosis inhibitor Ferrostatin-1 (**Figure 3F**, **Sup. 2D, E**). Consistently, the sensitivity to another ferroptosis inducer, the System Xc-inhibitor Erastin (XCTi) was significantly enhanced in CSFmax (**Figure 3G, Sup. 2F**).

**Figure 3:**
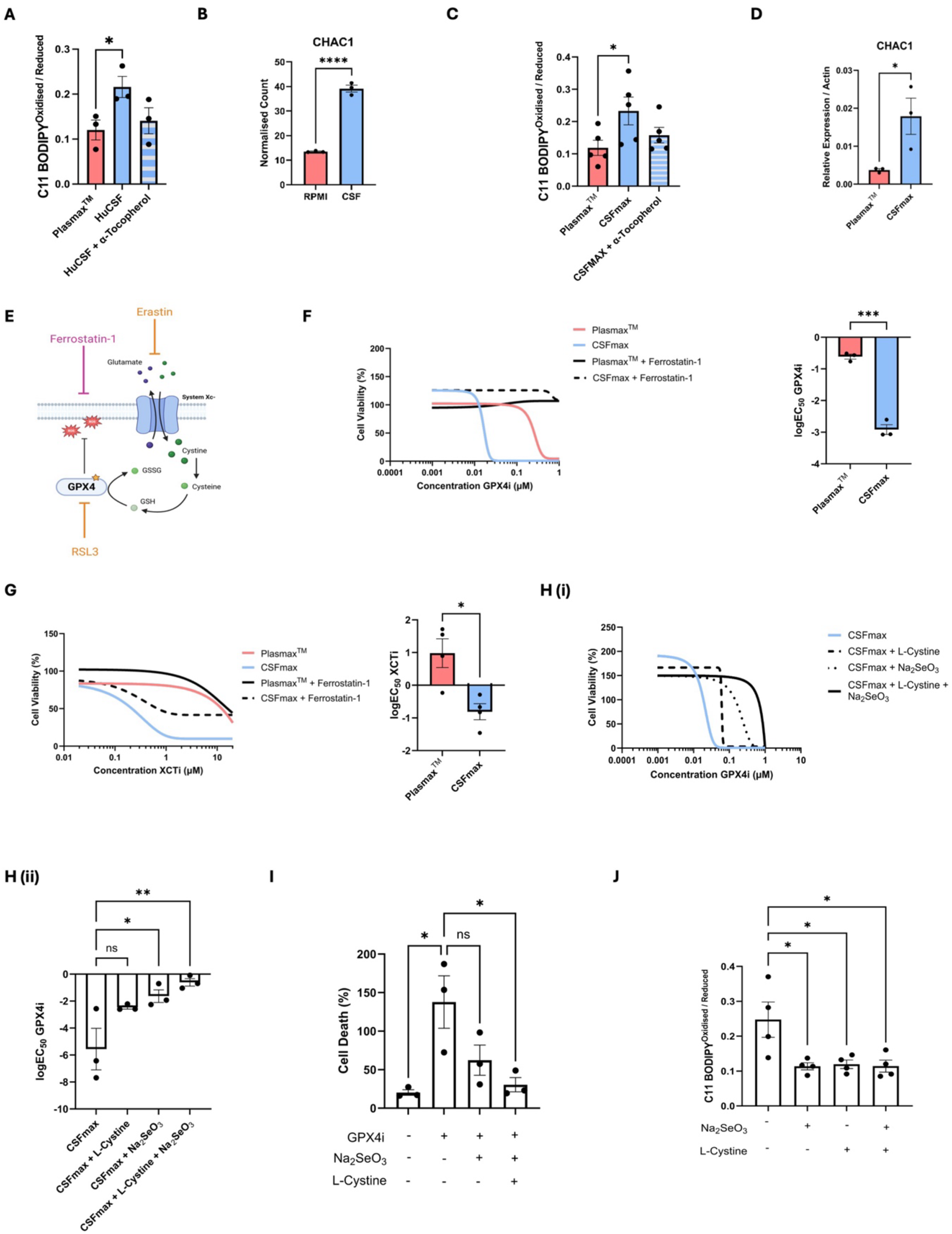
Cerebrospinal Fluid Primes ALL Cells for Ferroptosis. (**A**) Ratio of oxidised (510 nm) to reduced (591 nm) BODIPY 581/591 C11 in 697 cells incubated for 24 hours in Plasmax^TM^ or Patient CSF, with or without 20 µM α-tocopherol as indicated. N = 3 independent experiments. P-values calculated using unpaired two-tailed Student’s t-test. Error bars represent mean ± SEM. **(B)** *CHAC1* mRNA expression from publicly available RNA sequencing of ALL cells (REH) incubated in RPMI + 10% FBS or human CSF for 48 hours. P-values calculated using unpaired two-tailed Student’s t-test. (GSE274857). Error bars represent mean ± SEM. **(C)** Ratio of oxidised (510 nm) to reduced (591 nm) BODIPY 581/591 C11 in 697 cells incubated for 24 hours in Plasmax^TM^ or CSFmax, with or without 20 µM α-tocopherol. N = 5 independent experiments. P-values calculated using unpaired two-tailed Student’s t-test. Error bars represent mean ± SEM. **(D)** qPCR quantification of *CHAC1* mRNA expression, normalised to *ACTB* mRNA, in 697 cells treated for 48 hours in CSFmax or Plasmax^TM^. N = 3 independent experiments. P-values calculated using unpaired two-tailed Student’s t-test. Error bars represent mean ± SEM. **(E)** Schematic illustrating the mechanism of action of ferroptosis inducers RSL3 and Erastin (orange), and the ferroptosis inhibitor Ferrostatin-1 (pink). Created with BioRender.com. **(F)** Left Panel: Representative dose-response curves of 697 cells treated with RSL3 (GPX4i) in Plasmax^TM^, CSFmax, with or without 2 µM Ferrostatin-1 as indicated. Luminescence values for each concentration were normalised to vehicle-treated controls. N = 3 independent experiments. Right Panel: LogEC_50_ values in Plasmax vs CSFmax. Each point indicates the calculated logEC_50_ from independent experiments (n=3). P-values calculated using unpaired two-tailed Student’s t-test. Error bars represent mean ± SEM. **(G)** Left Panel: Dose-response curves of 697 cells treated with Erastin(XCTi) in Plasmax^TM^, CSFmax, with or without 2 µM Ferrostatin-1. Luminescence values for each concentration were normalised to vehicle-treated controls. N = 4 independent experiments. Right Panel: LogEC_50_ values in Plasmax vs CSFMax. Each point indicates the calculated logEC_50_ from independent experiments (n=4). P-values calculated using unpaired two-tailed Student’s t-test. Error bars represent mean ± SEM. **(H)** Upper Panel **(i)**: Dose-response curves of 697 cells treated with RSL3 in CSFmax with or without 30 nM Na_2_SeO_3_, with or without 65 µM L-Cystine as indicated. Luminescence values for each concentration were normalised to vehicle-treated controls. N = 3 independent experiments. Lower Panel **(ii)**: LogEC_50_ values. Each point indicates the calculated logEC_50_ from independent experiments (n=3).P-values determined by a one-way ANOVA with Dunnett’s multiple comparisons test. Error bars represent mean ± SEM. **(I)** Relative P.I. fluorescence of 697 cells treated with 75nM of RSL3 for 24 hours in CSFmax with or without 30 nM Na_2_SeO_3_, with or without 65 µM L-Cystine. Fluorescence normalised to lethal controls defined in the methods. N = 3 independent experiments. P-values determined by a one-way ANOVA with Tukey’s multiple comparisons test. Error bars represent mean ± SEM. **(J)** Ratio of oxidised (510 nm) to reduced (591 nm) BODIPY 581/591 C11 in 697 cells incubated for 48 hours in CSFmax, with or without 30 nM Na_2_SeO_3_ or 65 µM L-Cystine. N = 4 independent experiments. P-values determined by a one-way ANOVA with Dunnett’s multiple comparisons test relative to CSFmax alone. Error bars represent mean ± SEM. * ≤ 0.05 **** ≤ 0.0001

To identify the metabolic drivers of the enhanced ferroptotic sensitivity in CSF-like conditions we conducted nutrient rescue experiments. The supplementation of L-cystine and selenite at concentrations found in Plasmax™ rescued ferroptosis sensitivity to levels observed in Plasmax™-cultured cells (**Figure 3H-J**). These findings indicate that limited cystine and selenium availability are critical determinants of ferroptosis susceptibility in the CNS microenvironment.

### CSF-like Cystine Limiting Conditions are Synthetically Lethal with Selenocysteine Biosynthesis Interference

The 018Z BCP-ALL cell line has been used previously to model isolated CNS relapse *in vivo*^31^^32^. Interestingly, this line exhibits heightened ferroptosis resistance compared with other BCP-ALL cell lines (**Figure Sup. 3A**), suggesting intrinsic biological fitness under pro-ferroptotic conditions such as the CNS microenvironment. It has been shown that the ectopic expression of either FSP1 or GCH1 rescues the loss of GPX4 activity^33^^34^; as such, we measured *FSP1* and GCH1 expression in 018Z cells. *FSP1* showed similar expression in all cell lines (**Figure Sup. 3B).** However, in line with resistance to RSL3 treatment, we found that 018Z cells express the highest level of GCH1 amongst four BCP-ALL lines we tested (**Figure Sup. 3C**). GCH1 (GTP cyclohydrolase 1) is the rate-limiting enzyme in the biosynthesis of tetrahydrobiopterin (BH4), a potent lipophilic antioxidant that can inhibit ferroptosis^35^. Concordant with enhanced GCH1 function, 018Z cells demonstrated elevated levels of intracellular BH4 and its oxidised product BH2 (**Figure Sup. 3 D, E**). 018Z cells thus recapitulate the elevations of ferroptosis suppressing genes observed in CNS-relapse patient samples (**Figure 1D-G**). Therefore, in this stringent and clinically relevant model, we assessed the relevance of the anti-ferroptotic pathway centred around selenocysteine biosynthesis. CRISPR-Cas9 deletion of LRP8 had no effect on 018Z baseline growth (**Figure 4A, C**), nor sensitivity to FSP1 and GPX4 inhibition (**Figure Sup. 4A, B**). However, loss of LRP8 in 018Z sensitised to ferroptosis induced by cystine uptake inhibition (Erastin/XCTi) or cystine deprivation (**Figure 4B-E**). Cystine deprivation resulted in loss of GPX4 specifically in LRP8^KO^ cells (**Figure 4F**). Enhanced sensitivity to cystine uptake inhibition in LRP8^KO^ cells was phenocopied by alternative CRISPR-Cas9 deletion of SEPHS2 (**Figure 4G-J**). Collectively, these data indicate that in the CNS-tropic 018Z cells, despite the enhanced anti-ferroptotic activity of GCH1, the inhibition of selenium uptake and selenocysteine biosynthesis are synthetically lethal with cystine withdrawal.

**Figure 4:**
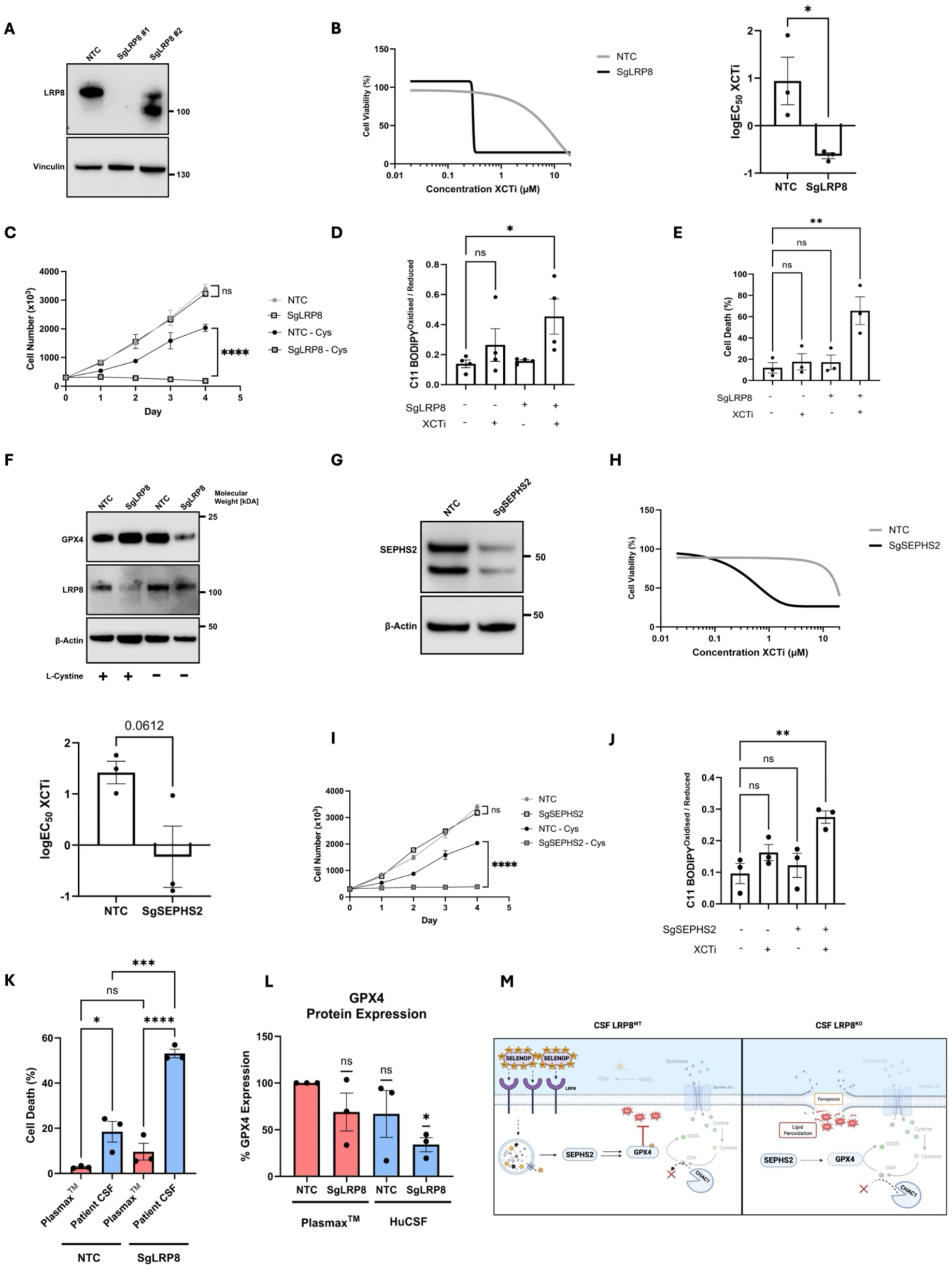
CSF-like Cystine Limiting Conditions are Synthetically Lethal with Selenocysteine Biosynthesis Interference. (**A**) Immunoblot images of LRP8 and Vinculin in 018Z transduced with NTC or with two LRP8 CRISPR guides (sg1 and sg2). **(B)** Left Panel: Dose-response curves of 018Z NTC or sgLRP8 #1 cells treated with Erastin in RPMI with 10%FBS. Luminescence values normalised to vehicle-treated controls. N = 3 independent experiments. Right Panel: logEC_50_ values for Erastin. Each point indicates the calculated logEC_50_ from independent experiments. P-values calculated using unpaired two-tailed Student’s t-test. Error bars represent mean ± SEM. **(C)** Cell Number of NTC or sgLRP8 #1 grown in media with or without 100 µM L-cystine over 96 hours. P-values determined by a one-way ANOVA with Tukey’s multiple comparisons test. N = 3 independent experiments. Error bars represent mean ± SEM. **(D)** Ratio of oxidised (510 nm) to reduced (591 nm) BODIPY 581/591 C11 in NTC or sgLRP8 #1 018Z cells incubated for 24 hours with or without 2.5 µM Erastin. N = 4 independent experiments. P-values determined by one-way ANOVA with Dunnett’s multiple comparisons test. Error bars represent mean ± SEM. **(E)** Relative P.I fluorescence of 018Z NTC or sgLRP8 #1 cells treated with 2.5µM Erastin for 48 hours. Fluorescence values were normalised to lethal controls described in the Methods. N = 3 independent experiments. P-values determined by a one-way ANOVA with Dunnett’s multiple comparisons test. Error bars represent mean ± SEM. **(F)** Immunoblot images of GPX4, LRP8 and β-Actin protein levels in sgLRP8 #1 or NTC cells cultured in media with or without 100 µM L-cystine for 48 hours. 1 experiment representative of 3 are shown. **(G)** Immunoblot image of SEPHS2 and β-Actin in 018Z cells transduced with NTC or SEPHS2 CRISPR guide. **(H)** Left Panel: Dose-response curves of 018Z NTC and sgSEPHS2 cells treated with Erastin in RPMI with 10% FBS. Luminescence values normalised to vehicle-treated controls. N = 3 independent experiments. Right Panel: logEC_50_ values for Erastin. Each point indicates the calculated logEC_50_ from independent experiments. P-values calculated using unpaired two-tailed Student’s t-test. Error bars represent mean ± SEM. **(I)** Cell number of NTC or sgSEPHS2 in media with or without 100 µM L-cystine over 96 hours. P-values determined by a one-way ANOVA with Tukey’s multiple comparisons test. N = 3 independent experiments. Error bars represent mean ± SEM. **(J)** Ratio of oxidised (510 nm) to reduced (591 nm) BODIPY 581/591 C11 in NTC or sgSEPHS2 018Z cells incubated for 24 hours with or without 2.5 µM Erastin. N = 3 independent experiments. P-values determined by two-way ANOVA with Dunnett’s multiple comparisons test. Error bars represent mean ± SEM. **(K)** Cell death in Plasmax^TM^ and human CSF. Flow cytometry quantification of DAPI-positive staining in NTC or sgLRP8 #1 cells cultured in Plasmax^TM^ or human CSF for 24 hours. P-values determined by two-way ANOVA with Tukey’s multiple comparisons test. N = 3 independent experiments. Error bars represent mean ± SEM. **(L)** Relative levels of GPX4 intracellular staining in sgLRP8 #1 or NTC 018Z cells after 48 hours in Plasmax^TM^ or human CSF. Data were normalised against Plasmax^TM^ treated NTC cells. N=3 independent experiments. P-value refers to a one-sample Wilcoxon signed-rank test. **(M)** Schematic representation of the proposed model whereby CSF induces compensatory upregulation of LRP8 to maintain GPX4 activity (*Left panel*) and that LRP8 targeting induces a synthetic lethality in CSF (*Right panel*). Error bars represent mean ± SEM. * ≤ 0.05 ** ≤ 0.01 **** ≤ 0.0001

To test the robustness of these observations across different contexts, we interrogated the results of an *in vivo* genome-wide CRISPR-Cas9 screen^36^ conducted to identify genes selectively required for survival of Jurkat T-ALL cell line in the CNS (**Figure Sup. 4C**). Several genes involved in the regulation of selenium and iron selectively affected the engraftment within the CNS compared to the bone marrow. For instance, NUBP2 (involved in regulating iron sulfur cluster formation), GPX1 (H_2_O_2_ scavenging selenoprotein), TXNRD1 (H_2_O_2_ scavenging selenoprotein) and PSTK (a key enzyme in selenocysteine biosynthesis), emerged as essential genes selectively in the CNS (**Figure Sup. 4D**). These data further support the notion that under conditions where GSH is perturbed, T-ALL rely on the antioxidant action of selenoproteins to survive. To cross-validate these observations in T-ALL, we transduced Jurkat cells with CRISPR-Cas9 guides targeting LRP8 (**Figure Sup. 4E**). The loss of LRP8 decreased GPX4 expression in Jurkat cells (**Figure Sup. 4E**) but was permissive for slow proliferation over several passages (**Figure Sup. 4F**). However, targeting LRP8 and cystine uptake or cystine withdrawal induced significant cell death, mirroring the synthetic lethality observed in 018Z cells (**Figure Sup. 4F-I**). Finally, we tested the effects of LRP8 deletion in 018Z cells cultured in patient CSF or Plasmax™. LRP8^KO^ cells had significantly higher levels of cell death in CSF (**Figure 4K**), accompanied by a marked reduction in GPX4 expression (**Figure 4L**). Together, these findings suggest that targeting LRP8 or selenocysteine biosynthesis is selectively lethal under cystine-limited conditions, an effect that is faithfully recapitulated in the cystine-scarce CSF (**Figure 4M**).

### Targeting Selenium Uptake and Utilisation Genes Inhibits CNS Leukaemia in vivo and Identifies Biomarkers of CNS Relapse

We next extended our findings using *in vivo* models of leukaemia. The 018Z cell line induces a highly aggressive leukaemia characterised by CNS tropism and rapid onset of hind limb paralysis within 3 weeks of injection^32^. We first assessed the impact of disrupting selenium utilisation on disease burden by genetically deleting SEPHS2 **(Figure 5A).** A significant reduction in both CNS and splenic tumour burden was observed with SEPHS2^KO^ cells, while their engraftment in the bone marrow remained unaffected (**Figure 5B**). To mechanistically dissect the effect of disrupted selenium uptake, we next engrafted LRP8^KO^ cells (**Figure 5C**). The tumour burden as well as lipid peroxidation levels were assessed. While LRP8 deletion did not affect leukaemic load in the bone marrow or spleen it selectively and significantly decreased CNS infiltration (**Figure 5D**). Notably, LRP8^KO^ cells exhibited increased lipid peroxides, particularly in cells colonising the CNS compartment (**Figure 5E**). These findings demonstrate that LRP8 suppresses lipid peroxidation, and that its loss is sufficient to compromise leukaemic cell survival in the CNS, despite having only a marginal effect on systemic disease. Given the pronounced reduction in CNS disease observed *in vivo* following interference with selenocysteine biosynthesis, we asked whether these genes could function as prognostic markers of CNS relapse in patient cohorts. We first assessed LRP8 expression in diagnostic bone marrow samples stratified by site of relapse. Patients who relapsed in the CNS had significantly higher *LRP8* mRNA levels compared with those who did not (**Figure 5F**), whereas no significant differences were observed when conducting the same analysis on patients who relapsed in the bone marrow (**Figure Sup. 5A**). Building on this observation, we explored the prognostic significance of the three genes highlighted in the current study — LRP8, SEPHS2 (whose knockout inhibited CNS engraftment) and PSTK (identified as a top CRISPR screen hit) using TARGET Phase II publicly available data^37^. In a multivariate analysis, upregulation of *LRP8* was the strongest risk factor for CNS relapse (OR 5.2, p=0.004), demonstrating greater impact than classical risk factors of CNS disease (age, MRD status and white blood cell counts (WBC)) (**Figure 5G**). Importantly, these associations were not recapitulated in analyses of bone marrow relapse (**Figure Sup. 5B**). To evaluate their combined effect, we derived a simple three-gene score based on the number of transcripts upregulated (Z-score ≥ 1). Fine-Gray cumulative incidence analysis demonstrated a stepwise increase in CNS relapse risk, patients with at least two upregulated genes had the highest relapse rates, while those with none had the lowest (**Figure 5H**). As expected, the same three-gene score showed no significant impact on bone marrow relapse (**Figure Sup. 5C**). Together, these findings support a model in which selenium uptake and utilisation pathways act as biological drivers of CNS leukaemia and suggest that their upregulation in leukaemic blasts may have prognostic relevance for risk of CNS relapse.

**Figure 5:**
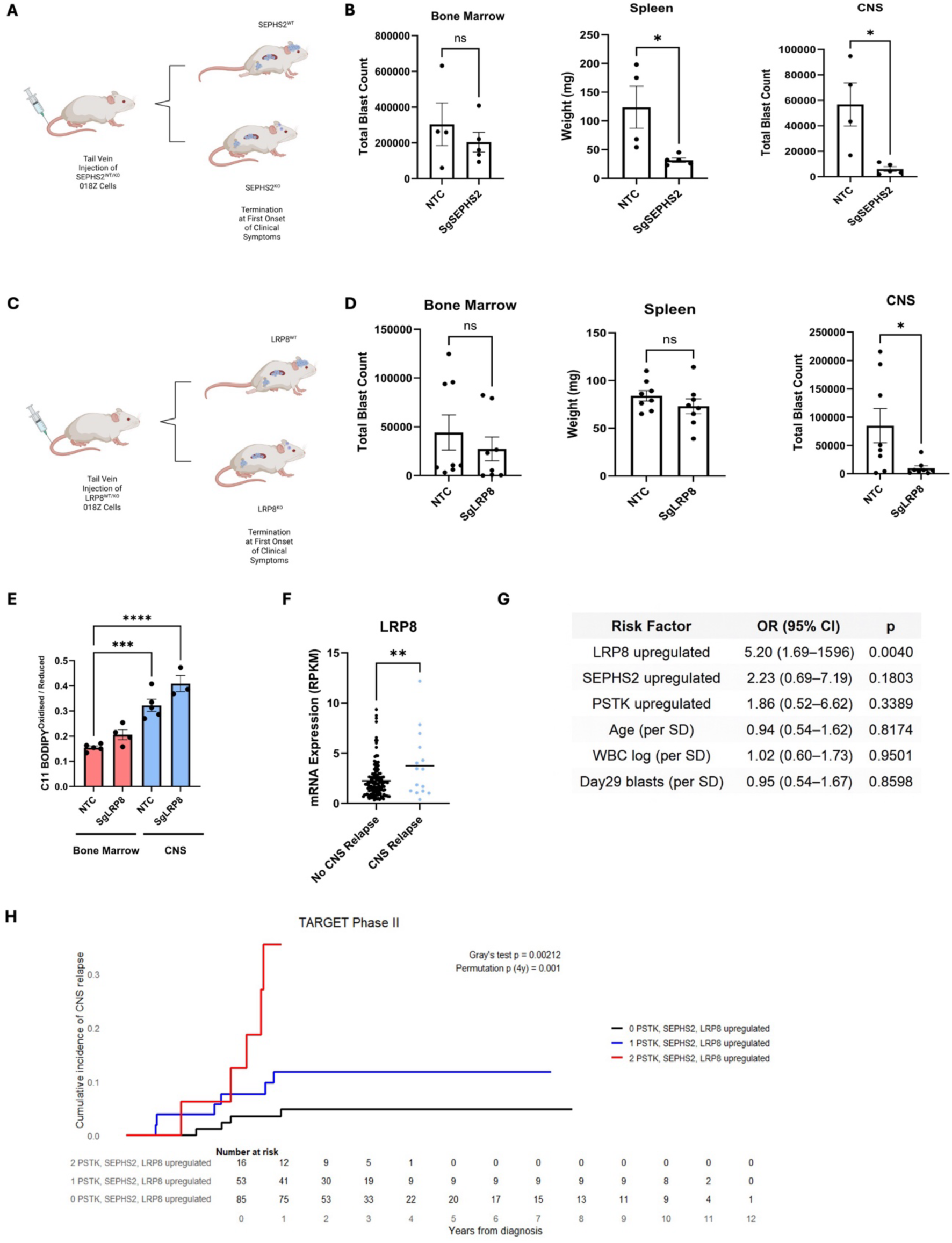
Targeting Selenium Uptake and Utilisation Genes Inhibits CNS Leukaemia In Vivo and Identifies Biomarkers of CNS Relapse. (**A**) Schematic of the xenotransplantation of NTC or sgSEPHS2 018Z cells. Created with BioRender.com. **(B)** Leukaemic burden assessed in the spleen by measuring its weight, or in the CNS and in the BM, by counting the total number of blasts. NSG mice were xenografted with sgSEPHS2 or NTC 018z cells. N= 4-5 mice per cohort as indicated by points. P-values calculated using unpaired two-tailed Student’s t-test. Error bars represent mean ± SEM. **(C)** Schematic of the xenotransplantation of NTC and sgLRP8 018Z cells. Created with BioRender.com. **(D)** Leukaemic burden assessed in the spleen by measuring its weight, or in the CNS and in the BM, by counting the total number of blasts. NSG mice xenografted with sgLRP8 #1 or NTC 018z cells. N= 8 mice per cohort as indicated by points. P-values calculated using unpaired two-tailed Student’s t-test. Error bars represent mean ± SEM. **(E)** Ratio of oxidised (510 nm) to reduced (591 nm) BODIPY 581/591 C11 in NTC or sgLRP8 #1 018Z cells obtained from BM or CNS. N=3-4 mice per experiment as indicated by points. P-values determined by a one-way ANOVA with Dunnett’s multiple comparisons test. Error bars represent mean ± SEM. **(F)** Diagnostic BM mRNA levels from TARGET Phase II patients (n=203) who relapsed in the CNS (n=23) compared to no-CNS relapse (n=180). P-values calculated using unpaired two-tailed Student’s t-test. Error bars represent mean ± SEM. **(G)** Cox Proportional Hazards model of risk of CNS relapse in this dataset with traditional risk factors (MRD positivity of >0.01% at day 29 of induction therapy, WCC > 50 × 10^9^/L, Age >10 years) and upregulation of LRP8, SEPHS2 or PSTK (Z-Score **≥** 1). **(H)** Fine–Gray analysis showing cumulative incidence of CNS-involving relapse in patients with upregulation of any one or two or more of LRP8, SEPHS2, or PSTK (gene expression z-score ≥ 1.0). * ≤ 0.05 *** ≤ 0.001 **** ≤ 0.0001

### Auranofin Serves as a Pharmacological Strategy to Target the Selenoproteome

To translate these findings into a novel therapeutic approach, we conducted a literature review to identify pharmacological compounds with two critical properties: (1) the ability to penetrate the blood-brain barrier (BBB) and (2) the capacity to disrupt selenoprotein biosynthesis. Auranofin (**Figure 6A**) emerged as a compelling candidate, consistently demonstrating both attributes across multiple studies^38^^39^^40^^41^^42^. The efficacy of Auranofin in inhibiting selenium incorporation into selenoproteins, GPX1 and GPX4, was first evaluated in 018Z and Jurkat cells treated with an excess of selenite (150 nM) to maximise selenoprotein expression, followed by incremental titration of Auranofin. Auranofin effectively suppressed selenoprotein biosynthesis within the nanomolar range, highlighting its therapeutic potential in this context (**Figure 6B, Figure Sup.6A**). To further explore the cellular response to Auranofin, we assessed its efficacy in CSFmax and Plasmax^TM^ media across a panel of different cell lines. Consistent with our hypothesis, cells grown in CSFmax were more sensitive to Auranofin (**Figure 6C-D, Figure Sup.6B-D**), an effect driven by cystine availability (**Figure 6E**). To assess Auranofin’s therapeutic potential *in vivo*, we engineered 018Z cells to express GFP and luciferase to monitor leukaemia progression. Luciferase signal was detectable in the bones of all mice by day 4 post-transplantation, and when the 10 mg/kg Auranofin was initiated (**Figure Sup.6E**). Mice displayed no clinical signs of Auranofin toxicity and weights remained stable throughout the experiment (**Figure Sup. 6F**). Mice were sacrificed upon emergence of hind limb paralysis in the vehicle-treated cohort or after 2 weeks of Auranofin treatment (Mon-Fri). Remarkably, Auranofin had no impact on splenic disease and minimal effects on bone marrow burden, but it significantly reduced leukaemia in the CNS (**Figure 6F**). To extend these findings, we employed a patient-derived xenograft (PDX) model in which PDX cells were injected via the tail vein, with weekly peripheral blood sampling conducted until the number of blasts exceeded 1% of host CD45 positive cells (**Figure 6G, Figure Sup. 6G**). At such point mice were randomised to receive either vehicle or Auranofin treatment. During a 2.5 week treatment period, no signs of drug-related toxicity were noted (**Figure Sup. 6H**). At the end of this period a predefined disease endpoint was reached and the experiment concluded. Consistent with the findings obtained with established cell lines, no significant differences were observed in spleen or bone marrow disease burden of Auranofin and vehicle treated mice (**Figure 6H**). Conversely, Auranofin significantly decreased the number of patient-derived leukaemic cells in the CNS (**Figure 6H**), where the levels of lipid peroxidation were selectively increased by the drug (**Figure 6I**). Furthermore, ALL blasts isolated from the spleen of vehicle– or Auranofin-treated mice revealed differential levels of selenoproteins (**Figure 6J**). SELENOP has two isoforms, detected by Western blot at 50kDa and 60kDa^43^^44^. The relative abundance of these isoforms has been shown to be regulated by plasma selenium levels in patients^45^. Auranofin eliminated the 50kDa isoform of SELENOP in leukaemic cells and significantly decreased the expression of GPX4 (**Figure 6J**). These results suggest that Auranofin inhibits selenocysteine biosynthesis *in vivo* and induces lipid peroxidation selectively in CSF resident ALL cells, thereby reducing disease burden within the CNS. Collectively, these findings support the discovery of a novel niche dependent metabolic vulnerability, whereby pharmacological inhibition of selenocysteine biosynthesis selectively targets CNS leukaemia.

**Figure 6:**
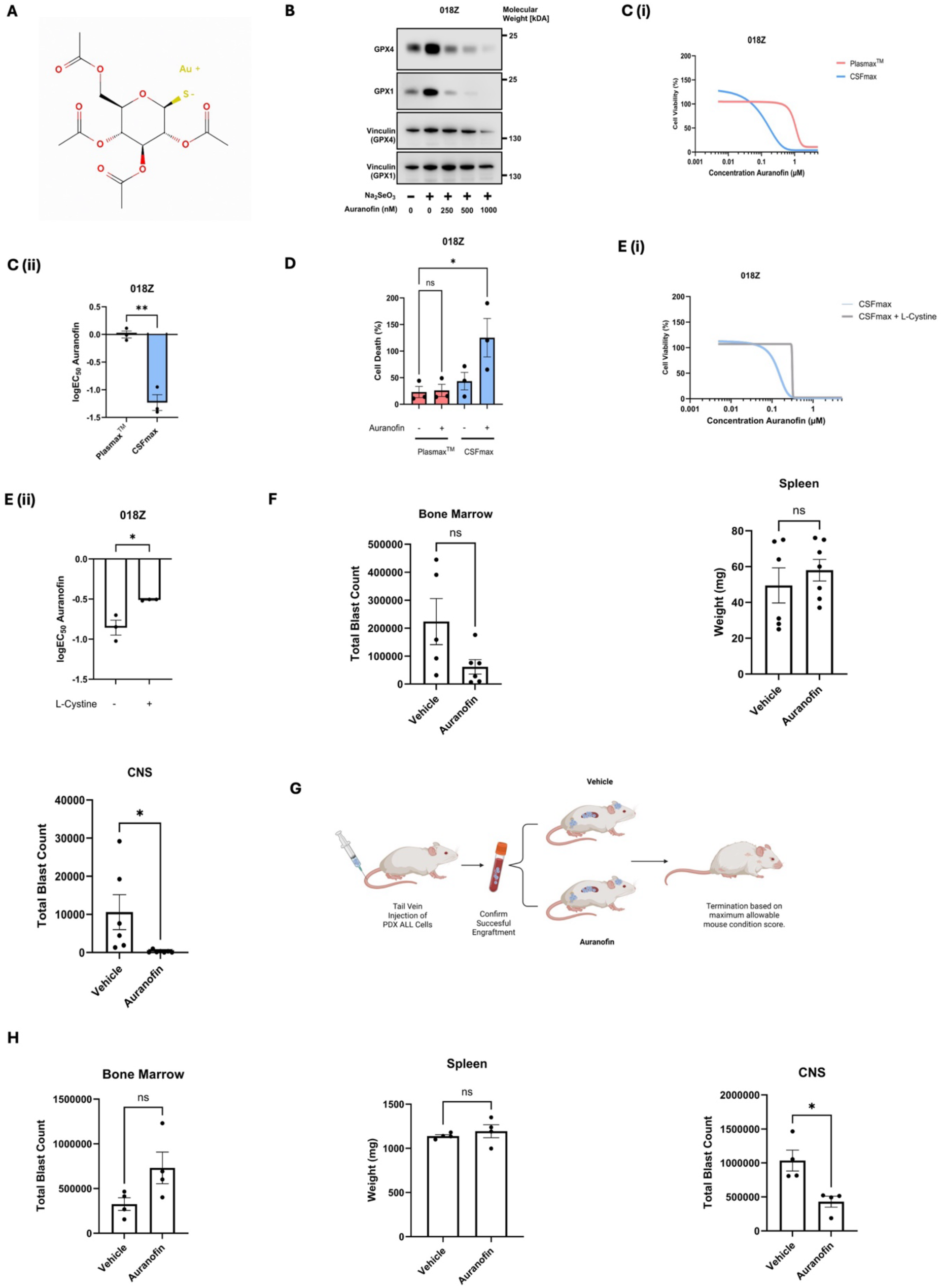
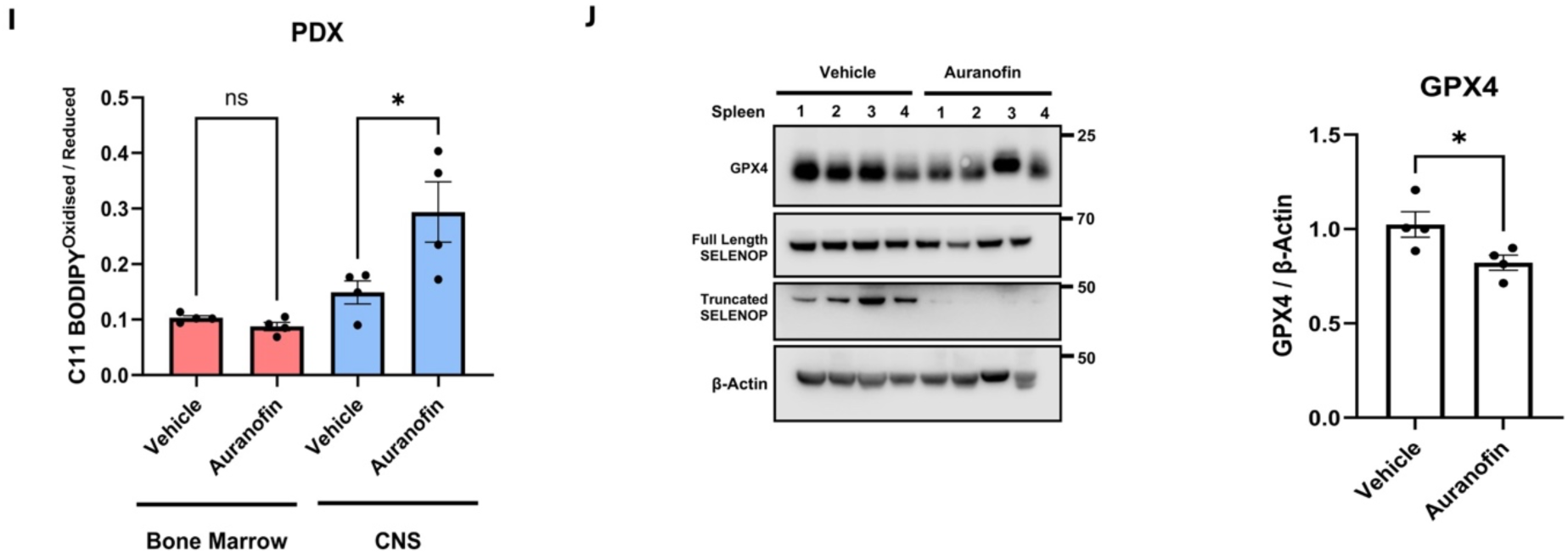
Auranofin as a Pharmacological Strategy to Target the Selenoproteome. (**A**) Auranofin chemical structure. Pubchem ID:16667669 **(B)** Immunoblot images of GPX4, GPX1 and Vinculin in 018Z cells treated with 150nM Na_2_SeO_3_ and with an increasing concentration of Auranofin for 72 hours as indicated. The results from 1 experiment representative of 3 are shown. **(C) Upper Panel:** Dose-response curves of 018Z, treated with Auranofin in CSFmax or Plasmax^TM^ as indicated. Luminescence values were normalised to vehicle-treated controls. Points indicate the mean of N = 3 independent experiments. **Lower Panel:** logEC_50_ values for Auranofin. Each point indicates the calculated logEC_50_ from independent experiments (n=3). P-values calculated using unpaired two-tailed Student’s t-test. Error bars represent mean ± SEM. **(D)** Relative P.I fluorescence of 018Z cells treated with 100nM Auranofin for 48 hours in Plasmax^TM^ or CSFmax. Fluorescence values were normalised to lethal controls described in Methods. P-values determined by a one-way ANOVA with Dunnett’s multiple comparisons test. Error bars represent mean ± SEM. **(E) Upper Panel:** Dose-response curves of 018Z treated with Auranofin in CSFmax with or without 65µM cystine. Luminescence values normalised to vehicle-treated controls. Points indicate the mean of N = 3 independent experiments. **Lower Panel:** logEC_50_ values for Auranofin. Each point indicates the calculated logEC_50_ from independent experiments (n=3). P-values calculated using unpaired two-tailed Student’s t-test. Error bars represent mean ± SEM. **(F)** Leukaemic burden assessed in the spleen by measuring its weight, or in the CNS and in the BM by counting the total number of blasts. NSG mice xenografted with human 018z ALL cell lines were treated with Auranofin (Mon-Fri). N= 6 or 7 mice per cohort as indicated by points. P-values calculated using unpaired two-tailed Student’s t-test. Error bars represent mean ± SEM. * ≤ 0.05 **(G)** Schematic of the patient-derived xenotransplantation and Auranofin treatment. Created with BioRender.com. **(H)** Leukaemic burden assessed in the spleen by measuring its weight, or in the CNS and in the BM by counting the total number of blasts. NSG mice xenografted with human 859I ALL PDX cells treated with Auranofin (Mon-Fri). P-values calculated using unpaired two-tailed Student’s t-test. Error bars represent mean ± SEM. **(I)** Ratio of oxidised (510 nm) to reduced (591 nm) BODIPY 581/591 C11 in Auranofin or vehicle treated cells obtained from BM or CNS of PDX xenograft at the time of sacrifice. N=4 mice. P-values determined by a two-way ANOVA with Tukey’s multiple comparison test. Error bars represent mean ± SEM. **(J)** Left Panel: Immunoblot images of GPX4, full length SELENOP, truncated SELENOP and β-Actin in samples from spleen of mice treated with vehicle or Auranofin as described in (**Figure 6G**) N=4 mice. Right Panel: Quantification of GPX4 protein levels. P-values calculated using unpaired two-tailed Student’s t-test. Error bars represent mean ± SEM. * ≤ 0.05.

## Discussion

Leukaemic blasts freely infiltrate the CNS without undergoing subclonal selection^46^. These data imply that relapse risk is dictated by the ability of leukaemic blasts to adapt and persist in the CNS microenvironment rather than selective entry^47^^48^. However, the mechanistic basis of this adaptation has until now been largely unknown. Here we show that low levels of cystine and thiols in CSF create a pro-oxidative environment and enhanced susceptibility to lipid peroxidation, revealing a niche-dependent metabolic vulnerability of ALL cells residing in the leptomeninges. These findings provide an underlying mechanism for our previous observation that SCD upregulation is a key determinant of successful CNS infiltration in ALL, a finding that appears conserved across CNS metastases of solid malignancies^8^. SCD promotes the synthesis of MUFAs, which are less prone to peroxidation and help maintain membrane integrity under oxidative stress. Thus, we demonstrate that successful adaptation to the CNS microenvironment by ALL cells necessitates both lipid desaturation and successful LRP8-mediated organic selenium scavenging to provide adequate antioxidant defence in the context of extremely low thiol and cystine levels available from the CSF.

Synthetic lethality between cystine deprivation and LRP8 loss has previously been observed in neuroblastoma cells with low expression of SLC7A11^25^. However, our findings extend this concept to a physiologically relevant *in vivo* context, where the microenvironment itself (rather than tumour-intrinsic factors) drives this vulnerability. This distinction is critical. The CSF enforces a ferroptosis-prone state not only on cells with preexisting metabolic liabilities, but potentially on all cells entering the CNS niche, regardless of their intrinsic redox capacity. Supporting this, we have consistently replicated our findings in both B and T-ALL cells across diverse genetic subtypes. Indeed, even in 018Z cells that overexpress GCH1 (part of an alternative ferroptosis resistance mechanism via BH4 synthesis)^34^, loss of LRP8 was synthetically lethal when combined with cystine depletion or exposure to human CSF. Interestingly, we found that LRP8 protein abundance is sensitive to thiol availability. How LRP8 detects thiol levels and responds by maintaining GPX4 expression through SELENOP uptake remains an open and important question for further investigation. Furthermore, the metabolic stress signature we observe in ALL cells appears conserved across multiple cancer types residing in the leptomeningeal niche. Models of leptomeningeal metastasis from breast, liver, and lung cancers similarly show upregulation of CHAC1, indicative of cystine deprivation and GSH depletion^49^. Moreover, we show that these solid tumours consistently upregulate Selenium Binding Protein 1 (SELENBP1), a selenium-binding protein previously implicated in anti-ferroptotic defence^25^, suggesting a common reliance on selenium metabolism in the CNS.

It remains unclear whether the combined targeting of GSH and selenocysteine biosynthesis reflects a phenotype driven exclusively by loss of GPX4 activity. Similar combinations – for example, SLC7A11-low neuroblastoma with LRP8^KO^—have previously been associated with partial ferroptosis rescue *in vivo*^25^, underscoring the complexity of this combinatorial cell death modality and highlighting the need for further mechanistic dissection. From a translational perspective, because direct GPX4 inhibition is likely to be toxic^50^, targeting selenocysteine – and specifically organic selenocysteine-mediated biosynthesis – may provide a more tractable therapeutic window. Moreover, as LRP8^KO^ and SELENOP^KO^ mice show no overt symptoms on a selenium-containing diet^51^^52^, the current study provides supporting evidence that interfering with organic selenium uptake could represent a novel and highly selective therapy for CNS ALL and, potentially, other leptomeningeal metastasis, bolstering the rationale for developing LRP8 inhibitors as therapeutic agents.

Our findings provide a strong rationale for further preclinical testing and evaluation of Auranofin in disease-relevant models. We show that Auranofin, an orally administered FDA-approved drug, can disrupt GPX4 via selenocysteine synthesis inhibition. To our knowledge, this is the first demonstration that Auranofin reduces selenoprotein abundance in cancer cells. We posit that prior observations made regarding anti-cancer effects of Auranofin^53^^54^^55^ and attributed to thioredoxin reductase (*TXNRD1*) inhibition^56^ could be also explained, at least in part, by its impact on global selenoprotein translation. Furthermore, given its well-characterised safety profile and long-standing clinical use, Auranofin presents a promising candidate for repurposing in early-phase trials for CNS relapse. Notably, Auranofin has demonstrated low toxicity in paediatric populations^57^, which is a critical consideration in developing new therapies for children. Its oral route of administration offers a major practical and clinical advantage over current intrathecal therapies, which require repeated general anaesthesia – a procedure associated with high cost, logistical burden, and potential neurotoxicity with cumulative exposure^58^. Although we did not directly assess CNS penetrance in our models, prior studies demonstrate BBB penetration of Auranofin^41^^42^, and our data suggest selective efficacy in CNS disease. Further preclinical pharmacokinetic and dynamic studies will be required to de-risk translation.

In conclusion, we identify a previously unrecognised metabolic vulnerability in leptomeningeal cancers arising from the unique metabolic composition of CSF. CNS leukaemic cells rely on LRP8-mediated selenium uptake and SEPHS2-dependent selenocysteine biosynthesis to maintain redox balance and ferroptosis-associated defences under cystine-limited conditions. This adaptive loop constitutes a liability that can be pharmacologically exploited, as shown with Auranofin, and may be further targeted through inhibition of LRP8 or other components of the selenocysteine pathway. To our knowledge, this selenium dependence has not previously been described in cancer, highlighting a niche-specific therapeutic opportunity for CNS leukaemia and potentially other leptomeningeal metastases.

## Methods

### Publicly Available Patient mRNA Sequencing Data

Patient and PDX data were downloaded from the GEO database (https://www.ncbi.nlm.nih.gov/geo/), Refs. <u>GSE60926</u> and GSE271998. Data were assessed for normalisation and differential expression analysis was conducted using DESEQ2. Other data sets GSE271998, GSE83132 and RNA sequencing previously published by our laboratory^7^ were obtained, normalised and either DESEQ2 was conducted, or *CHAC1* mRNA levels were obtained. Clinically annotated RNA sequencing and microarray data from the TARGET Phase II trial were retrieved using the cBio Cancer Genomics Portal^59^.

### Cell culture

Cell lines used throughout this work are detailed in **Supplementary Table 2**. Cultures of Acute Lymphoblastic Leukaemia (ALL) cells CCRF-CEM (ATCC, CCL119), Jurkat (ATCC) 697 (DSMZ, #ACC42), KOPN8 (DSMZ, #ACC552) SEM (ATCC), REH (ATCC, CRL-8286) and 018Z were maintained in RPMI-1640 (Thermo Fisher Scientific, #31870025) supplemented with 10%FBS (Thermo Fisher Scientific, # 10270106), L-glutamine 2mM (Thermo Fisher Scientific, #25030024), 1% penicillin/streptomycin (Thermo Fisher Scientific, #15140122) solution. The 018Z cell line was a kind gift from Professor Doctor Luder Meyer. Cells were cultured at 37°C in ambient O_2_ (∼21%) in a 5% CO_2_ incubator and were regularly monitored for mycoplasma contamination and authenticated using Promega GenePrint 10 Kit (Promega, #B9510).

### Preparation of CSFmax

CSFmax represents an in-house produced physiological media designed to mimic closely the nutrient and biochemical composition of human CSF. In brief, metabolite concentrations in CSF from adult individual were sourced from the Human Metabolome database^20^ and Wishart et al.,^60^ and the concentrations of electrolytes from clinical studies^61^^62^^63^^7,64^. Individual metabolites and salts were reconstituted from powder into suitable solvent (e.g. water). These solutions were further diluted in a buffered solution containing sodium bicarbonate, phenol red, Plasmax^TM^ cell culture media (CancerTools.org, #156371) and BME vitamin mix (Sigma, #B6891). Finally, media was supplemented with 3.75% delipidated dialysed FBS and 1.5% dialysed FBS (Capricorn Scientific, # FBS-DIA-12A). FBS was delipidated using fumed silica precipitation^9^.

### Patient Samples and Ethical Approval

Diagnostic ALL live cells were sourced from the Bloodwise (now VIVO) Childhood Leukaemia Cell Bank; cerebrospinal fluid (CSF) samples from ALL patients were obtained via the West of Scotland CSF Biobank; normal control CSF was obtained from the Glasgow Neuroimmunology Biobank; and plasma samples were acquired from the NHS Greater Glasgow & Clyde Bio-repository. Individuals providing ‘healthy subject’ CSF were enrolled prior to receiving any formal diagnosis and presented with a range of neurological symptoms. At the time of sample collection, all CSF tests were normal, and no specific neurological condition had been confirmed, therefore these individuals were therefore classified as ‘healthy subjects’ for the purposes of this study. Use of all biobanked human samples for this project was approved by the West of Scotland Research Ethics Service (reference 09/S0703/77), and all research was conducted in accordance with the Declaration of Helsinki.

### Cloning and CRISPR-based gene editing

The following gRNA sequences: Non-Targeting Control (NTC): 5’-GTAGCGAACGTGTCCGGCGT-3’, LRP8 g1 5’-GGCCACTGCATCCACGAACGG-3’, LRP8 g2 5’ – GCTGCTTAGACCACAGCGACG –3’, *SEPHS2*: 5’-GAGGGACGGCAGTGACCGG-3’ cloned into lentiCRISPRv2 vector (Addgene, Plasmid #52961) using BsmBI restriction sites. For lentivirus production, 2×10^6^ HEK293T cells were transfected with 5 µg lentiCRISPRv2 plasmid (NTC or on-target), 1µg pVSV-G (Addgene, #12259) and 3µg psPAX2 (Addgene, #12260). Six hours after transfection the medium (DMEM, 10% FBS, 2mM L-glutamine without antibiotics) was replaced, and cells incubated for 18 hours before the medium was harvested for viral infection of recipient cells. The recipient 018Z or Jurkat cells were cultured for 48 hours with lentivirus-containing medium supplemented with 8 µg/ml Polybrene (Merck, # TR-1003) and then for additional 48 hours with fresh medium, before undergoing a 4-day selection in medium supplemented with 1.5μg/mL puromycin (Thermo Fisher Scientific, # A1113903). After transduction, LRP8– and SEPHS2-knock-out pools were cultured in RPMI with 10% FBS and 2mM L-glutamine, 1% penicillin/streptomycin solution.

### Xenografting

All animal experiments were approved by Institutional Ethical Review Process Committees and were performed under UK Home Office license (PPL PP5432144). *JAX NOD.Cg-Prkdc^scid^Il2rg^tm1Wjl^/SzJ* (NSG; Charles River Laboratories, Harlow, United Kingdom) mice were kept in sterile isolators with autoclaved food, bedding, and water. Experimental groups consisted of individual male and female animals. Appropriate age– and sex-matched control groups were used per experiment, xenografts of lentiCRISPRv2 NTC cells in genetic knockout experiments and the vehicle drug solvent in Auranofin-treated animals were used as controls. Exact n of animals is reported in each figure legend. No *a priori* exclusion criteria were used. Where disease burden was assessed by IVIS imaging or blood sampling prior to Auranofin treatment, mice were assigned to experimental groups to ensure that the disease burden was comparable between groups, therefore randomisation was not used. Experimental conduct, outcome assessment and data analysis were not blinded. Mice were weighed and monitored using scoring charts for body condition, behavioural and clinical signs of leukaemic engraftment. Xenotransplants were performed in 6-10 week old NSG mice by tail vein injection of cells resuspended in sterile PBS. Cells were mycoplasma tested prior to injection. At the end of experiment, mice were sacrificed using CO_2_ and cells were collected from bone marrow, spleen and leptomeninges as previously described^65^. For protein extraction, leukaemic cells were purified using a Ficoll gradient (Lymphoprep, Stemcell) following the supplier instructions.

For assessment of disease burden, spleens were weighed, and bone marrow and leptomeningeal cells were analysed to check human engraftment by FACS using AttuneNxT excluding mouse CD45 positive (1:1000, Thermo Fisher Scientific; 11-0451-82) and counting cells positive for human CD19 (1:1000, Biolegend; 302224). Dead cells were removed using propidium iodide (P.I) exclusion (1µg/ml, Thermo Fisher Scientific; P1304MP). Patient derived cells were initially xenografted (PDX) in cohort of NSG mice to expand them and obtain enough cells for secondary experimental xenografts, where 1 × 10^6^ cells / mouse were injected in another cohort to perform the experiments. Flow cytometry data were analysed using FlowJo software (BD Biosciences). Auranofin (MedChemExpress, #HY-B1123) or vehicle (PBS) was administered to mice by an i.p. injection at 10 mg per kg body weight 5 days/week (Mon-Fri). Tissues were collected after 10 and 13 administrations of Auranofin or vehicle in 018Z and PDX experiments respectively, unless mice were sacrificed sooner due to early signs of hind-limb paralysis. For the 018Z model, one vehicle-treated and one auranofin-treated mouse were excluded from bone marrow analysis due to abnormally high leukaemic burden values which fell outside the distribution of remaining data points. These outlier values were identified using the ROUT method (Q = 1%) implemented in GraphPad Prism 9.0. The result of the experiment remained unchanged following removal.

### Bioluminescence Imaging

018Z Cells were transduced with PHAGE PGK-GFP-IRES-LUC-W (Addgene #46793), a stable luciferase expressing construct and sorted based on GFP expression. To guide commencement of Auranofin treatment in 018Z experiment, leukaemia burden was assessed after 4 days using bioluminescence imaging by injecting mice subcutaneously with 200 μL (15 mg/ml) D-Luciferin (Promega, # E1601), anesthetising them with 2–3% isoflurane, and imaging them on an IVIS Spectrum (PerkinElmer). Representative image is presented in **Figure 6J**.

### Immunoblotting

Cells were lysed in radioimmunoprecipitation assay (RIPA) buffer. Protein was quantified using the Bradford assay (Thermo Fisher Scientific, #23200). Lysates were incubated in NuPAGE™ LDS Sample Buffer (4X) (Thermo Fisher Scientific, #NP0007) at 95⁰C for 3 min and loaded onto a SDS-polyacrylamide gel (4-12%, Invitrogen NuPAGE, #NP0336BOX). After size separation, proteins were transferred onto re-activated polyvinylidene fluoride (PVDF) membranes, which were subsequently blocked in 5% defatted milk in Tris-buffered saline, 0.1% Tween-20 (TBST) for 1 h room temperature (RT). Primary antibodies were diluted in a solution of 5% bovine serum albumin (BSA) in TBST and incubated overnight at 4°C. Membranes were washed and then incubated at RT for 2 h with horseradish peroxidase-labelled secondary (1:5,000, anti-rabbit #7074, or anti-mouse #7076 Cell Signaling) diluted in a solution of 5% milk in TBST. The following antibodies were used against β-actin (1:10,000; 66009-1-Ig, Proteintech), Vinculin (1:5000; 700062, Invitrogen), GPX4 (1:1,000; ab125066, Abcam), LRP8 (1:500; ab108208, Abcam), GPX1 (1:1,000; ab22604, Abcam), SEPHS2 (1:1000; 14109-1-AP, Proteintech), SELENOP (1:1000; PA5-28281, Invitrogen), GCH1 (1:1000; H00002643-M01, Abnova).

### Ellman’s Assay

Ellman’s assay (Merck, #D8130) measurement for thiol groups was performed as previously described^66^ using paired plasma and CSF samples taken from 2 patients. Thiol concentration was determined as described^66^. Pathlength was assumed to be 0.62 cm for 200 µL in a flat-bottom 96-well plate, consistent with Tecan Infinite® pathlength correction documentation.

### Liquid Chromatography–Mass Spectrometry (LC-MS)

Metabolomics analysis was carried out as previously described^67^. Briefly, chromatographic separation of metabolite extracts was performed using a ZIC-pHILIC column (SeQuant; 150 mm × 2.1 mm, 5 µm; Merck) along with a ZIC-pHILIC guard column (SeQuant; 20 mm × 2.1 mm; Merck) integrated with a Vanquish HPLC system (Thermo Fisher Scientific). A gradient method was utilised, using 20 mM ammonium carbonate (pH 9.2, containing 0.1 % v/v ammonia and 5 µM InfinityLab deactivator (Agilent)) as mobile phase A and 100% acetonitrile as mobile phase B. The elution began with 20% A for two minutes, followed by a linear increase to 80% A over 15 minutes, concluding with a re-equilibration step returning to 20% phase A. The column oven was maintained at 45 °C, with a flow rate to 200 µL min⁻¹. Metabolite analysis and identification were conducted using a Q Exactive Plus Orbitrap mass spectrometer (Thermo Fisher Scientific) equipped with electrospray ionisation. The instrument operated in polarity switching mode at a resolution of 70,000 at 200 m/z, allowing the detections of both positively and negatively charged ions over a mass range of 75 to 1,000 m/z. The automatic gain control (AGC) target was set to 1 × 10^6^, with a maximal injection time (IT) of 250 ms. Data analysis was undertaken in Skyline (version 23.1.0.455)^68^. Identification was accomplished by matching accurate mass and retention time of observed peaks to an in-house library generated using metabolite standards (mass tolerance of 5 ppm and retention time tolerance of 0.5 min).

### Metabolite Extraction from Patient CSF

CSF samples stored at –80°C were thawed on ice. Once thawed, 10 μL of CSF were diluted 1:20 with extraction solvent (50% methanol / 30% acetonitrile / 20% deionised water) and mixed thoroughly by vortexing for 30 seconds. The samples were then centrifuged at 16,000 x g for 10 minutes at 4°C. Before each extraction, 5 μL were taken from every CSF sample and pooled together separately in a 2 mL centrifuge tube to create a pooled quality control (QC) sample used for normalisation and for calibration.

### Intracellular GSH Measurement

2 × 10^6^ SEM cells were transplanted in 4 female NSG immunodeficient mice. When mice reached the overall severity threshold, the BM and CNS were harvested, human CD19^+^ cells were isolated, and metabolites were extracted and analysed through LC-MS. To measure intracellular levels of metabolites, huCD19^+^ cells were collected using Lymphoprep density gradient according to manufacturer’s instructions. After ensuring through flow cytometry, that the purification of human CD19^+^ cells harvested is above 95%, metabolites were extracted and analysed. Firstly, cell concentration was measured using the CASY automated cell counter (Roche). Cell concentration counts were used to normalise the volume of extraction solvent to achieve a concentration of 1 × 10^6^ cells/mL in all experimental conditions. Cells were washed twice with ice-cold phosphate buffered saline (PBS) solution (Cat. #14190094) with pulse centrifugation at 12,000 g for 15 sec at 4 °C. Following this, cells were suspended with the appropriate volume ice-cold extraction solvent (50:30:20, v/v/v acetonitrile/methanol/water) and were vortexed for 30 sec and incubated at 4 °C for 5 min. Cells were then centrifuged at 16,000 g for 10 min at 4 °C and the supernatant was transferred to liquid chromatography-mass spectrometry (LC-MS) glass vials. LC-MS glass vials were stored at –80 °C until measurements were performed. Equal volumes of extract were analysed by LC–MS.

### Intracellular BH2/BH4

ALL cells (SEM, 018Z) were cultured in CSFmax for 2 passages. At the 3rd passage, cells were seeded in fresh media at 1.7×10^5^/mL and cultured for 7 hours. Polar metabolites were extracted with slight modifications to protocol from^34^. Cell concentration counts were used to normalise the volume of extraction solvent to achieve a concentration of 1×10^6^ cells/mL in all experimental conditions. Cells were washed once quickly in ice-cold PBS followed by addition of ice-cold extraction buffer consisting of methanol:acetonitrile:water (v/v, 5:3:2) with freshly prepared 250μM ascorbic acid and 0.5% DTT. Next, samples were subjected to 16,000g at 4°C for 10min and supernatant collected for LCMS analysis. Equal volumes of extract were analysed by LC–MS.

### Quantitative (q) PCR

Cells were seeded in 6 well plate at density and timeframe as reported in figure legends. After seeding, the cells were washed with ice-cold PBS and pelleted at 4⁰C at 10,000g for 30s. RNA was isolated from cell pellets following the kit manufacturer’s protocol (Qiagen RNeasy, # 74104). RNA concentration was measured with the NanoDrop 2000 Spectrophotometer (Thermo Fisher Scientific). 500 ng RNA was used for cDNA synthesis (SuperScript VILO MasterMix, # 11755-050). 2ng cDNA and 50 nmol of each primer were used in each quantitative (q) real-time polymerase chain reaction qPCR was performed with Fast SYBR Green Master Mix (Applied Biosystems Fast SYBR Green Master Mix, #4385612) on a thermocycler (Bio-Rad). Cycling: 50 °C 2 min; 95 °C 2 min; 40 cycles of 95 °C 15 s, 58.5 °C 15 s, 72 °C 60 s. Primer sequences, ACTB: FWD (5’-3’) GGCATGGGTCAGAAGGATT RVS (5’-3’) ACATGATCTGGGTCATCTTCTC, CHAC1: FWD (5’-3’) GAACCCTGGTTACCTGGGC RVS (5’-3’) CGCAGCAAGTATTCAAGGTTGT, AIFM2 (FSP1): FWD (5’-3’) AGACAGGGTTCGCCAAAAAGA RVS (5’-3’) CAGGTCTATCCCCACTACTAGC

### Propidium Iodide (P.I.) Cell Death Assay

Cell death was assessed using propidium iodide (P.I) (Thermo Fisher Scientific, P1304MP) uptake measured on a TECAN plate reader. Cells were seeded at 2.5 × 10⁵ cells per ml in 6 well plate. At indicated experimental timepoint, P.I was added directly to each well at a final concentration of 1 µg/mL, and plates were incubated at 37°C for 15–30 minutes. Fluorescence was measured using a TECAN Infinite® plate reader (Ex: 535 nm, Em: 636 nm). Background fluorescence from P.I-only controls was subtracted, and values were normalised to cells treated with 10µM ABT737 (ApexBio, #A8193) and 2.5µM S68645 (Chemgood, #C-1370) for 6 hours serving as a lethal control.

### DAPI Viability Assay

NTC or LRP8^KO^ 018z cells were seeded at 3×10⁴ cells/well in 96-well plates in either Plasmax or patient CSF and cultured at 37 °C/5% CO₂ for 24hours. At endpoint, wells were gently mixed to resuspend all cells and the full contents were transferred to tubes, washed once with PBS, and resuspended in 100–200 µL PBS containing 1 μg/ml 4′,6-diamidino-2-phenylindole (DAPI) (Thermo Fisher Scientific, # D1306). Samples were acquired on an Attune NxT using the VL1 channel (≥10,000 events/sample). Unstained controls were used to set gates. Data were analysed by sequential gating for debris exclusion (FSC/SSC), singlets (FSC-A vs FSC-H), and live/dead discrimination, with DAPI⁺ events scored as non-viable and DAPI⁻ as viable; viability was reported as % DAPI^+^ of singlets for each condition.

### Viability Assay

Cell viability was assessed using the Cell-Titer Glo luminescence assay. Cells were seeded on 96-well plates at 3 × 10^4^ cells/well in 200μl culture media (RPMI 10%FBS, RPMI without cystine and methionine (Merck, # R7513), (RPMI 10% delipidated FBS), Plasmax (CancerTools, #156371) or CSFmax as indicated in figure legend) for 24 h 1S3R-RSL3 (Selleckchem, #S8155), 48 h (Erastin (Selleckchem, # S7242) and Auranofin) and 72 h (iFSP1 (Medchemexpress, #HY-136057)), (SW203668 (Cambridge Biosciences, #GC17354)). Compounds were dissolved in DMSO (Merck, # 472301) and further diluted in PBS (Thermo Fisher Scientific, # 14190094). For dose–response assays, a two-fold, 12-point dilution series of each compound was prepared. 100 µL of drug dilution was added to 100 µL of cell suspension per well (final volume 200 µL). Each assay was performed in technical triplicate (wells) which were averaged. Figures show the mean of n = 3 independent experiments. logEC_50_ values were obtained by non-linear regression (four-parameter logistic) on base-10 log-transformed concentrations. Cell death inhibitors were used at the following concentrations 10μM Q-VD-Oph (AdooQ Bioscience, #A14915-25), 20μM α-tocopherol (Merck, #258024), 2µM Ferrostatin (Merck, #SML0583), 65µM L Cystine (Medchemexpress, #HY-N0294) or 30nM Sodium selenite (Merck, # S5261) and supplemented to cell culture at the same time of the cytotoxic agents. Cells were then incubated for the indicated time points. The assays as described above were quantified using the ATP-based bioluminescence Cell-Titer Glo (Promega, #G7572). To determine cell viability, the Cell Titer Glo reagent and cell culture medium from each well were mixed 1:1 and incubated at room temperature for 10 min. Luminescence was assessed on a Promega GloMax Microplate Reader. Data were normalised against vehicle containing control well.

### Proliferation Assay

018Z or Jurkat cells were seeded at a density of 0.5 × 10⁶ cells/ml in a 6-well plate. Indicated treatments were added to three wells per condition. Cell numbers were measured at 24h intervals using the CASY automated cell counter (Roche).

### C11-BODIPY Lipid Peroxidation Assay

Lipid peroxidation was quantified using the C11-BODIPY 581/591 probe (Thermo Fisher Scientific, # D3861). Cells were seeded in 6-well plates at a density of 0.5 × 10⁶ cells per well in appropriate media indicated in figure legend. Following the treatments, cells were incubated with 1.5 µM C11-BODIPY in appropriate culture media for 30 minutes at 37°C in the dark. After staining, cells were washed with cold PBS, resuspended in FACS buffer (PBS + 2.5 % FBS), and immediately analysed by flow cytometry. The oxidised (510–550 nm) and reduced (565–605 nm) forms of the dye were detected using standard flow cytometry channels (i.e., FITC and PE). The ratio of the signal from the two fluorescence wavelengths was used as a relative quantitation for the lipid peroxidation. For the assessment of lipid peroxidation ex vivo, mice were sacrificed at the indicated time points, and bone marrow, spleen, CNS/meningeal compartment were harvested as previously described^46^. Single-cell suspensions were prepared by mechanical dissociation and filtering through 40 µm strainers. Cells were washed and resuspended in PBS containing 1.5 µM C11-BODIPY and incubated for 30 minutes at 37°C in the dark. After staining, cells were washed with cold PBS and stained with anti-human CD19 or anti-human CD3 (T-ALL PDX). Lipid peroxidation was quantified based on the oxidised/reduced C11-BODIPY fluorescence ratio within the leukaemic human CD19⁺ or CD3^+^ population. Flow cytometry data were analysed using FlowJo software (BD Biosciences).

### GPX4 Intracellular Staining

For assessment of intracellular GPX4 levels in human CSF, 3 × 10^4^ cells/well 018Z NTC or LRP8^KO^ cells were plated in 200 µL of patient CSF or Plasmax^TM^ in 96 well plate for 48 hours. Cells were collected by centrifugation (500 × g, 5 min), washed with PBS, and fixed in 4% paraformaldehyde for 10 minutes at room temperature. After fixation, cells were washed and permeabilised using Flow Cytometry Permeabilisation/Wash Buffer I containing 0.2% Triton X-100. For intracellular staining, cells were incubated with a GPX4 primary antibody (1:350 dilution, no. ab125066, Abcam) for 15 minutes, followed by a single wash. Cells were subsequently incubated with an Alexa Fluor-conjugated secondary antibody (1:1000, A-11012, Thermo Fisher Scientific) for 30 minutes at room temperature, protected from light. Following a final wash, cells were resuspended in PBS and analysed using flow cytometry. Flow cytometry data were analysed using FlowJo software (BD Biosciences).

### Genome-wide CRISPR Screen

Jurkat human T-ALL cells were transduced with lentiviruses containing the Brunello human CRISPR knockout pooled library which encompasses 76,411 gRNAs targeting 19,114 genes. Human Brunello CRISPR knockout pooled library was a gift from David Root and John Doench (Addgene #73179; http://n2t.net/addgene:73179; RRID:Addgene_73179)^69^. A total of ∼150 million cells were transduced at a multiplicity of infection (MOI) of 0.3 to achieve coverage of approximately 500 cells per sgRNA. Cells were selected for puromycin (Merck #TR-1003, 1 µg/mL) for 72h. Approximately 4 × 10^7^ puromycin resistant cells were expanded for a week. During this time cells were passaged and maintained at ∼40 million cells. Collected cells (∼40 million cells) were divided equally and xenotransplanted in 13 NSG mice (Charles River Laboratories, Harlow, United Kingdom). BM and CNS resident cells were retrieved upon signs of leukaemic disease burden as described above. A total of ∼20 million and ∼10 million cells were retrieved from BM and CNS respectively. Genomic DNA from these cells was isolated with the QIAamp DNA Blood Maxi Kit (Qiagen #51192) followed by PCR amplification using Illumina adapters to identify the gRNA representation in each cell.

### Statistical Analysis

Each experiment was performed in ≥ 2 independent replicates. N refers to independent replicates, which are biologically distinct samples or experimental repeats conducted on different days using separately prepared cell cultures, or in different xenografted animals, or using patient-derived material from different individuals. All t-tests and ANOVA analyses were performed using GraphPad Prism version 9.0 and are described in figure legend. Appropriate post-hoc multiple comparisons tests (e.g. Dunnett’s or Tukey’s, as specified in figure legends) were applied following ANOVA. No a priori power calculations were conducted to determine the n number.

### Author Contributions

NG, SaT, ST and CH conceived the study. NG, RC, SL, EH, AM, VA, JM, KD, DO’C performed the scientific investigation. NG, RC, AM, JJS, AC, EH, ES, AHU, DO’C and DS analysed and visualised the data. NG, RC, SL, TA, EH, AM, JJS, ES, AHU, SB, VH, AC, DS, JVV, SaT, DO’C, MM, ST, and CH contributed to methodology development. CH and ST supervised the work. NG, SaT and CH wrote the original draft, and all authors critically reviewed and revised the manuscript for important intellectual content.

### ICMJE Authorship Compliance Statement

All listed authors meet the International Committee of Medical Journal Editors (ICMJE) criteria for authorship.

### Ethical Approval

This study was conducted in accordance with the principles of the Declaration of Helsinki. Ethical approval for the study was obtained from the West of Scotland Research Ethics Committee (WoSREC ID:09/S0703/77) under the protocol titled *“Understanding the Causes of CNS Leukaemia”*.

## Acknowledgements

This work was generously supported by a Cancer Research UK (CRUK) Programme Foundation Award to CH (DRCPFA-Nov21\100001), a Children’s Cancer and Leukaemia Group (CCLG) award to CH (CCLGA 2020 24), a CRUK Programme Award to ST (DRCNPG-Jun22\100011) a CRUK Clinical Academic Training Programme (TRACC) fellowship awarded to NG (SEBCATP-2022/100004) and CRUK Scotland Centre funding (CTRQQR-2021\100006). The authors gratefully acknowledge the Cancer Sciences Flow Cytometry Facility, University of Glasgow, for their support and assistance in this work. Samples and data used in this study were provided by VIVO Biobank, supported by Cancer Research UK & Blood Cancer UK (Grant no. CRCPSC-Dec21\100003). The authors thank John Goodfellow, Yasar Yousafzai, Antony Cousins and Saeeda Bhatti for assistance with CSF biobanking and especially the patients and their families who provided samples for this research. For the purpose of open access, the authors have applied a Creative Commons Attribution (CC BY) licence to any Author Accepted Manuscript version arising from this submission.

## Data Availability Statement

Source data are available from the corresponding authors upon reasonable request.

## Code Availability Statement

All custom scripts used for data analysis and figure generation in this study are available upon reasonable request from the corresponding author. The code was implemented in R (v4.3) for statistical analyses, RNA-seq processing, and visualisation. Standard packages (e.g., DESeq2, ggplot2) were utilised as specified in the Methods section. No proprietary or commercially restricted software was used.

## Declaration of Interests Statement

SaT is the inventor of Plasmax^TM^ cell culture medium. CH, SaT, EK and VH are named inventors on a patent application for CSFmax cell culture medium (filing date 09/05/2025, patent application number 2507177). All other authors declare no competing financial interests or personal relationships that could have appeared to influence the work reported in this study.

## Extended data/supplemental figures

**Supplemental Table 1:** Results of literature review on selenium speciation in mammalian CSF versus plasma.

| Species | Plasma /CSF | Quantification | Units | n | Reference | Means of Detection |
| --- | --- | --- | --- | --- | --- | --- |
| Total Se | Plasma | 59.67 | ug/L | 24 (Paired) | Solovyev, Berthele and Michalke, 2013 | ICP-DRC-MS |
| SELENOP | Plasma | 5.19 | ug/L | 24 (Paired) | Solovyev, Berthele and Michalke, 2013 | ICP-DRC-MS |
| Total Se | CSF | 2.371 | ug/L | 24 (Paired) | Solovyev, Berthele and Michalke, 2013 | ICP-DRC-MS |
| SELENOP | CSF | 0.474 | ug/L | 24 (Paired) | Solovyev, Berthele and Michalke, 2013 | ICP-DRC-MS |
| Total Se | CSF | 2745 | ng/L | 68 | Maass et al., 2020 | ICP-sf-MS |
| SELENOP | CSF | 2020 | ng/L | 68 | Maass et al., 2020 | ICP-sf-MS |
| Total Se | CSF | 2.3 | ug/L | 42 | Mandrioli et al., 2017 | ICP-sf-MS |
| SELENOP | CSF | 1.923 | ug/L | 42 | Mandrioli et al., 2017 | ICP-sf-MS |
| Total Se | Plasma | 82 | ng/g | 5 | M. A. Garcia-Sevillano, T. Garcia-Barrera and J. L. Gomez-Ariza, 2014 | IDA-ICP-ORS-MS |
| SELENOP | Plasma | 52 | ng/g | 5 | M. A. Garcia-Sevillano, T. Garcia-Barrera and J. L. Gomez-Ariza, 2014 | IDA-ICP-ORS-MS |
| Total Se | Plasma | 93.7 | ug Kg -1 | 23 | Del Castillo Busto et al., 2024 | ICP-MS |
| SELENOP | Plasma | 65.3 | ug Kg -1 | 23 | Del Castillo Busto et al., 2024 | ICP-MS |

**Supplemental Table 2:** Results of literature review on cystine reference values in patient CSF versus plasma.

| Plasma / CSF | Reference Concentration | Units | Reference |
| --- | --- | --- | --- |
| Plasma | 17-98 | μM | Allegri G, et al. 2010 |
| Plasma | 8.0 - 68.0 | μM | BC Children's Hospital Biochemical Genetics Lab. |
| Plasma | 62.9 +/- 27.8 | μM | Psychogios N, et al. 2011 |
| CSF | 0-1.5 | μM | BC Children's Hospital Biochemical Genetics Lab. |
| CSF | 0.29 +/- 0.27 | μM | Engelborghs S, et al. 2003 |
| CSF | 0.1 +/- 0.1 | μM | Geigy Scientific Tables, 8th Rev edition, pp. 165-177. |

**Supplemental Table 3:** Details of cell lines and patient derived xenograft (PDX) samples.

| Cell Line | Cytogenetic Code | Age at sampling | Sex | Immunophenotype | Resource Identification Initiative/source |
| --- | --- | --- | --- | --- | --- |
| 697 | t(1;19)(q23;p13) | 12Y | M | B cell precursor | 697 (RRID:CVCL_0079) |
| 018Z | 47, XY, +8, del(9)(p13) | Adolescent | M | B cell precursor | https://doi.org/10.1182/blood.V114.22.1630.1630 |
| CCRF-CEM | t(8;9)(p11;p24)x2 der(9)del(9)(p21-22)del(9)(q11q13-21)x2 | 4y | F | T cell | CCRF-CEM (RRID:CVCL_0207) |
| Jurkat | add(2)(p21)/del(2)(p23)x2 | 14Y | M | T cell | Jurkat (RRID:CVCL_0065) |
| KOPN8 | t(11;19)(q23;p13) KMT2A-MLLT1 | 3M | F | B cell precursor | KOPN-8 (RRID:CVCL_1866) |
| SEM | t(4;11) KMT2A-AFF1 | 5Y | F | B cell precursor | SEM (RRID:CVCL_0095) |
| 859I (PDX) | t(1;19) | 12Y | F | B cell precursor | Supplied by VIVO Biobank (https://vivobiobank.org) |

**Supplemental Figure 1:**
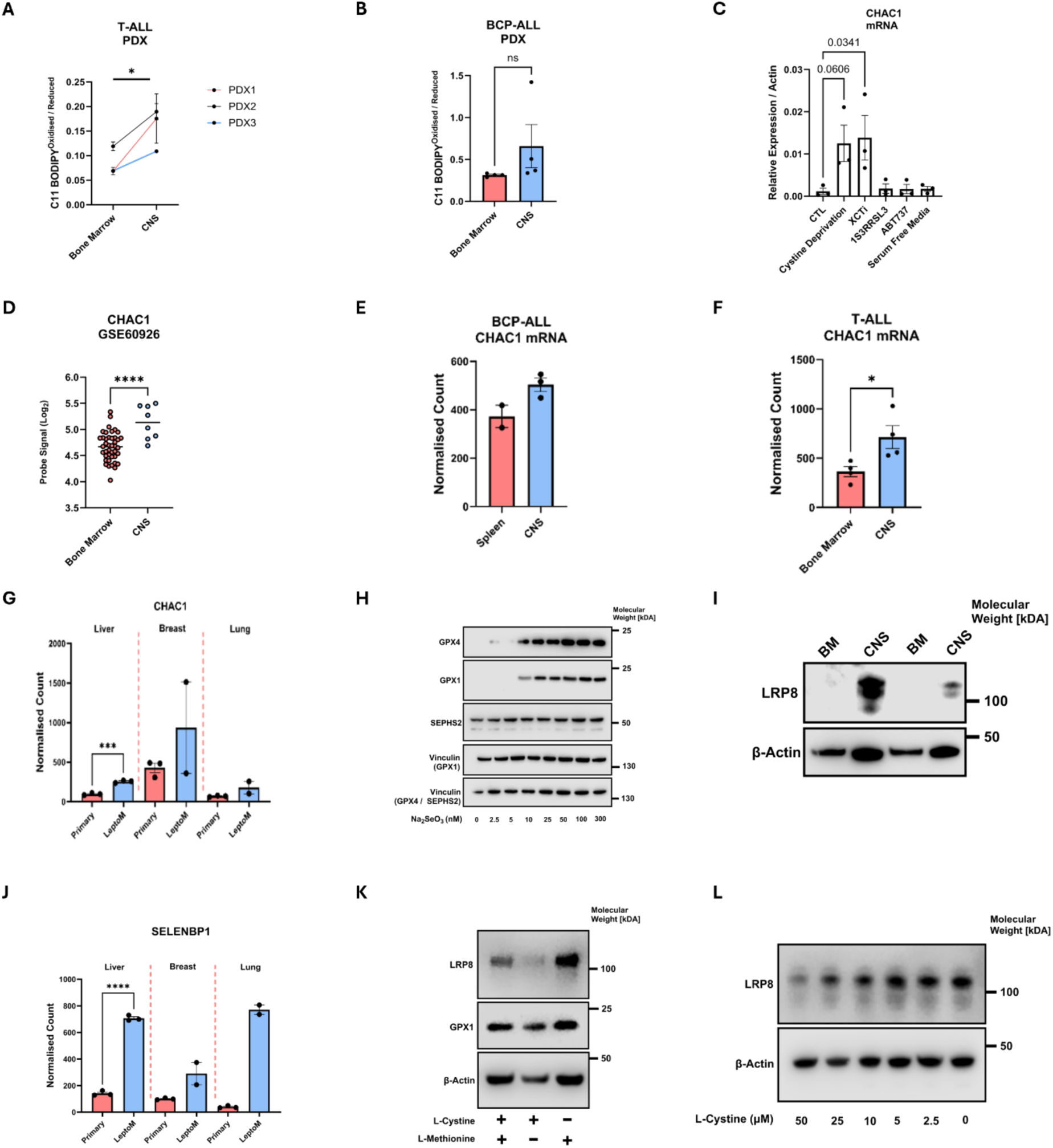
CNS-resident Cancer Cells Display Disrupted Glutathione Metabolism. **(A)** Ratio of oxidised (510 nm) to reduced (591 nm) BODIPY 581/591 C11 (lipid peroxidation sensor) in BM and CNS of 3 T-ALL PDX’s, PTMM (PDX1), OWL4-SP1 (PDX2) and 58A-SA1 (PDX3). P-values refer to a two-way ANOVA comparing lipid peroxidation in BM and CNS and PDX line. No significant differences were observed between PDX models. **(B)** Ratio of oxidised (510 nm) to reduced (591 nm) BODIPY 581/591 C11 (lipid peroxidation sensor) in BM and CNS from BCP-ALL 859I PDX cells. **(C)** qPCR quantification of *CHAC1* mRNA expression, normalised to *ACTB* mRNA abundance in 018Z cells treated for 24 hours in RPMI without cystine, or with 5 µM Erastin, 250 nM RSL3, 250 nM ABT737, or in serum-free RPMI. n = 3 independent experiments. P-values refer to a one-way ANOVA test with Dunnett’s multiple comparisons test relative to the untreated control. Error bars represent mean ± SEM. **(D)** *CHAC1* expression levels in leukaemic BM samples (n = 42) and CNS relapse samples (n = 8) from GSE60926. Each dot represents an individual sample (red = BM, blue = CNS). P-values calculated using an unpaired two-tailed Student’s t-test. **(E)** *CHAC1* mRNA expression in REH cells retrieved from the spleen or CNS of xenografted mice (n = 2 or 3, as indicated by points). Error bars represent mean ± SEM. **(F)** *CHAC1* mRNA expression in CCRF T-ALL cells retrieved from the BM or CNS of xenografted mice (n = 4). P-values calculated using unpaired two-tailed Student’s t-test. Error bars represent mean ± SEM. **(G)** CHAC1 expression levels from GSE83132 assessed in primary tumour or leptomeningeal metastasis of PC9 (lung) HCC1954 (liver) or MDAMB231 (breast) cancer cell lines. P-values calculated using unpaired two-tailed Student’s t-test. Error bars represent mean ± SEM. **(H)** Immunoblot images of GPX4, GPX1, SEPHS2 (GPX1 membrane) and Vinculin in 018Z cells cultured for 48 hours in serum free media with incremental increases in Na_2_SeO_3_. Results from 1 of 3 independent experiments are shown. **(I)** Immunoblot image of LRP8 and β-Actin in 859I PDX BCP-ALL cells isolated from the CNS and BM of two additional xenografted mice at clinical endpoint. These results were obtained from the biological replicates (n = 2) not shown in the main figure **(Figure 2H**), where data from 3/5 representative mice were presented. Blots confirm consistent protein expression trends across all five animals. **(J)** SELENBP1 expression levels from GSE83132 assessed in primary tumour or leptomeningeal metastasis of PC9 (lung) HCC1954 (liver) or MDAMB231 (breast) cancer cell lines. P-values calculated using unpaired two-tailed Student’s t-test. Error bars represent mean ± SEM. **(K)** Immunoblot images of LRP8, GPX1 and β-Actin in 018Z cells cultured in cystine-methionine free media treated with 100 µM methionine or 100 µM cystine for 48 hours as indicated. Results from 1 of 2 independent experiments are shown. **(L)** Immunoblot image of LRP8 and β-Actin in 018Z BCP-ALL cells cultured in cystine-free media supplemented with indicated concentrations of L-Cystine over 48 hours. Results from 1 of 3 independent experiments are shown. * ≤ 0.05 *** ≤ 0.001 **** ≤ 0.0001

**Supplemental Figure 2:**
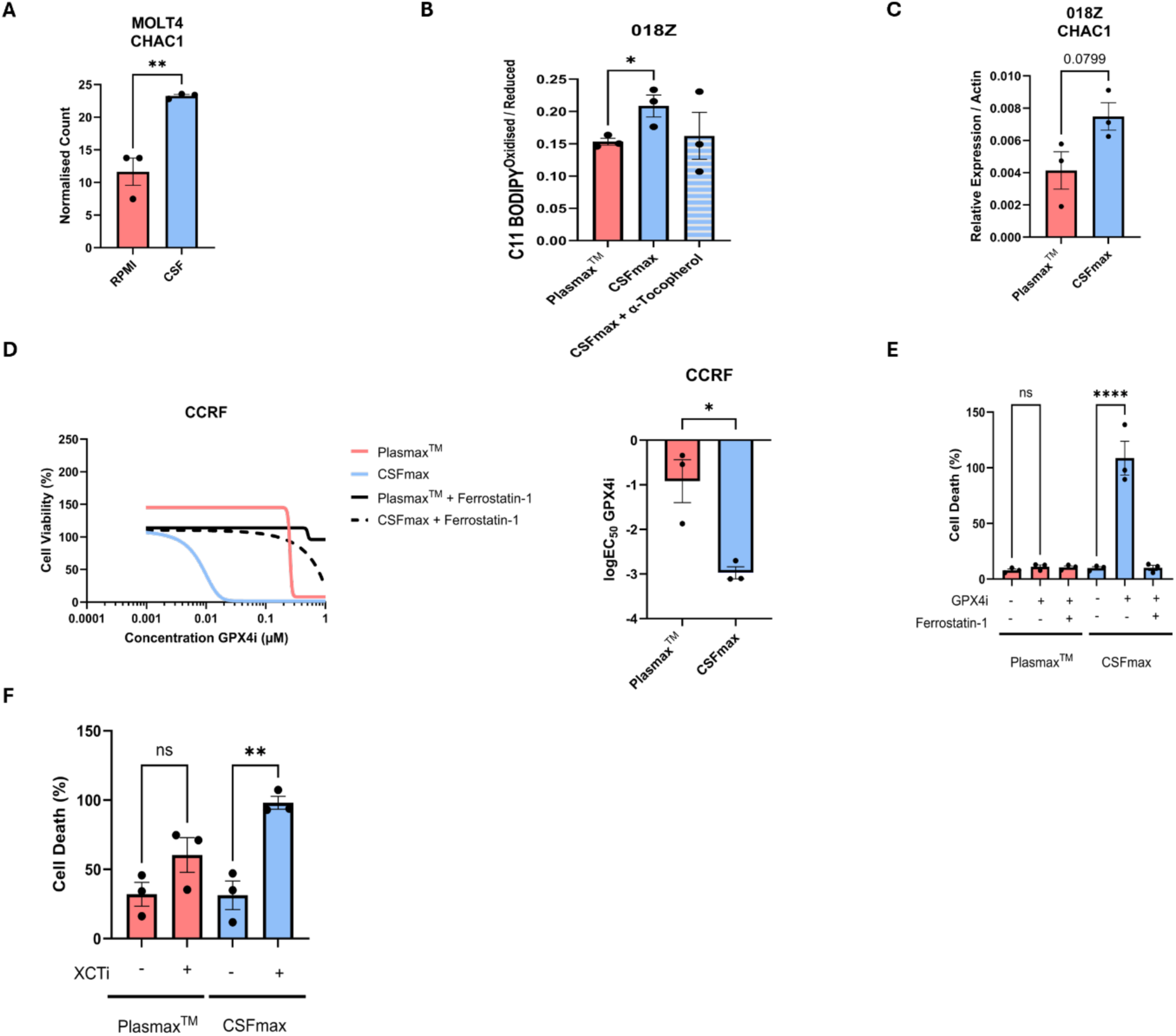
Cerebrospinal Fluid Primes ALL Cells for Ferroptosis. **(A)** *CHAC1* mRNA expression from publicly available RNA sequencing of ALL cells (MOLT-4) incubated in RPMI + 10% FBS or human CSF for 48 hours (GSE274857). P-values calculated using unpaired two-tailed Student’s t-test. Error bars represent mean ± SEM. **(B)** Ratio of oxidised (510 nm) to reduced (591 nm) BODIPY 581/591 C11 in 018Z cells incubated for 24 hours in Plasmax^TM^ or CSFmax, with or without 20 µM α-tocopherol. N = 3 independent experiments. P-values calculated using unpaired two-tailed Student’s t-test. Error bars represent mean ± SEM. **(C)** qPCR quantification of *CHAC1* mRNA expression, normalised to *ACTB* mRNA, in 018Z cells treated for 48 hours in CSFmax or Plasmax^TM^. N = 3 independent experiments. P-values calculated using unpaired two-tailed Student’s t-test. Error bars represent mean ± SEM. **(D)** Left Panel: Representative dose-response curves of CCRF cells treated with RSL3 in Plasmax^TM^, CSFmax, with or without 2 µM Ferrostatin-1 as indicated. Luminescence values for each concentration were normalised to vehicle-treated controls. N = 3 independent experiments. Right Panel: LogEC_50_ values for RSL3 Each point indicates the calculated logEC_50_ from independent experiments. P-values calculated using unpaired two-tailed Student’s t-test. Error bars represent mean ± SEM. **(E)** Relative P.I. fluorescence of 697 cells treated with 75 nM RSL3 for 24 hours in Plasmax^TM^ or CSFmax, with or without 2 µM Ferrostatin-1. Fluorescence values were normalised to lethal control described in the Methods. N = 3 independent experiments. P-values determined by a one-way ANOVA with Tukey’s multiple comparisons test. Bars represent mean ± SEM. **(F)** Relative P.I. fluorescence of 697 cells treated with 1.5µM Erastin for 48 hours in Plasmax^TM^ or CSFmax. Fluorescence values were normalised to lethal control described in the Methods. N = 3 independent experiments. P-values determined by a one-way ANOVA with Tukey’s multiple comparisons test. Error bars represent mean ± SEM. * ≤ 0.05 ** ≤ 0.01 ≤ 0.0001

**Supplemental Figure 3:**
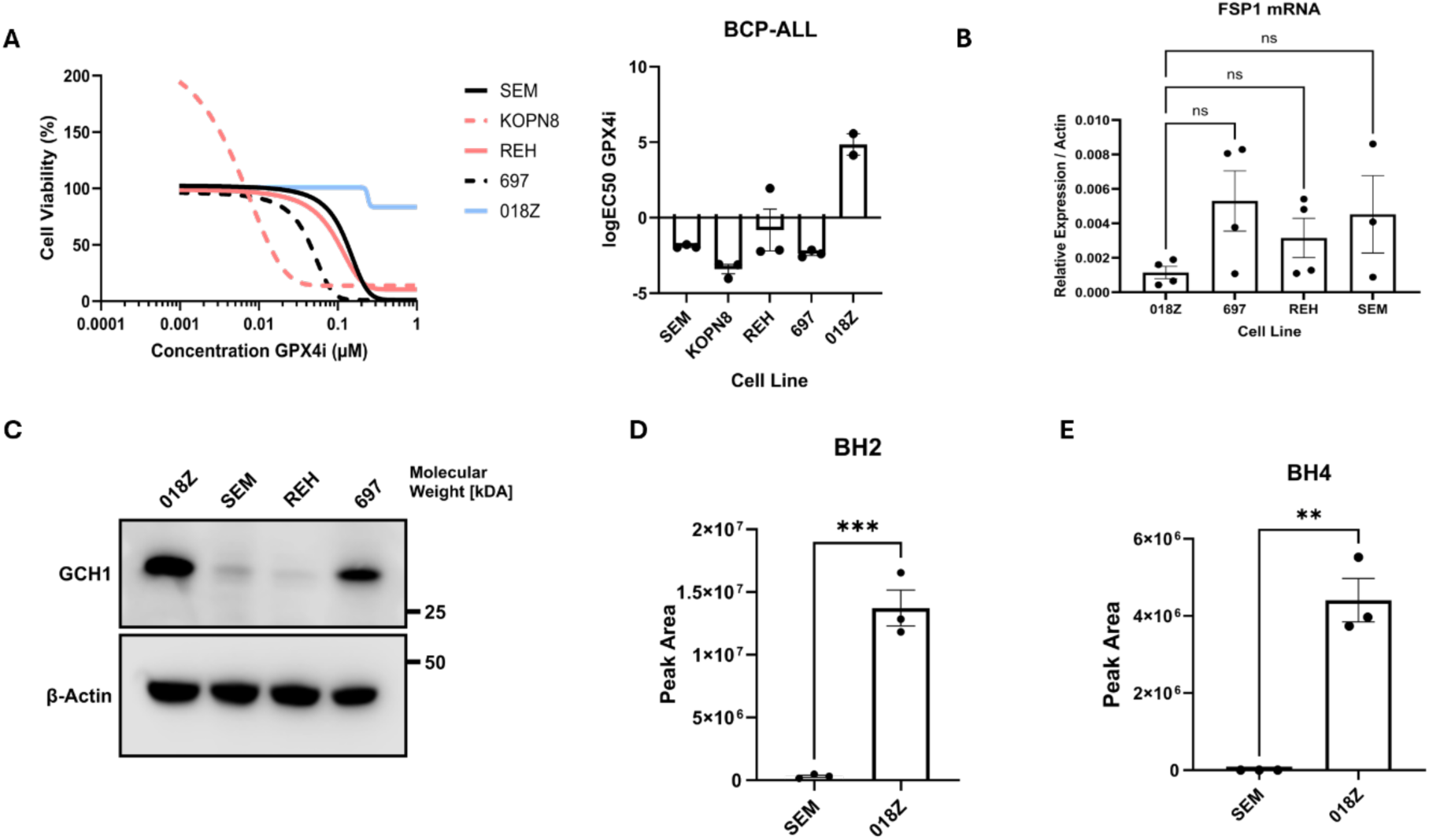
Isolated CNS Model Cell Line 018Z are Intrinsically Resistant to Ferroptosis Owing to Enhanced BH4 Synthesis. **(A)** Left: Dose–response curves of SEM, KOPN8, 697, REH, and 018Z cells treated with RSL3 in RPMI supplemented with 10% FBS. Luminescence values were normalised to vehicle controls. Right: logEC_50_ values, with each point representing an independent experiment (N = 3; 1 018Z curve could not be fitted due to lack of cytotoxicity). Data are shown as mean ± SEM. **(B)** qPCR quantification of *FSP1* mRNA expression in SEM, 697, REH, 018Z cells, normalised to ACTB. N = 4 independent experiments as indicated by the data points. P-values determined by a one-way ANOVA with Tukey’s multiple comparisons test. Error bars represent mean ± SEM. **(C)** Immunoblot images of GCH1 and β-Actin from 018Z, SEM, REH and 697 BCP ALL cells. 1 representative of 3 independent experiments. **(D)** Intracellular levels of BH2 in SEM and 018Z cells. Peak area normalised to cell count and shown as arbitrary units per 10⁶ cells. N = 3 independent experiments. P-values calculated using unpaired two-tailed Student’s t-test. **(E)** Intracellular levels of BH4 in SEM and 018Z cell line. Peak area normalised to cell count and shown as arbitrary units per 10⁶ cells. N = 3 independent experiments. P-values calculated using unpaired two-tailed Student’s t-test. Error bars represent mean ± SEM. ** ≤ 0.01 *** ≤ 0.001.

**Supplemental Figure 4:**
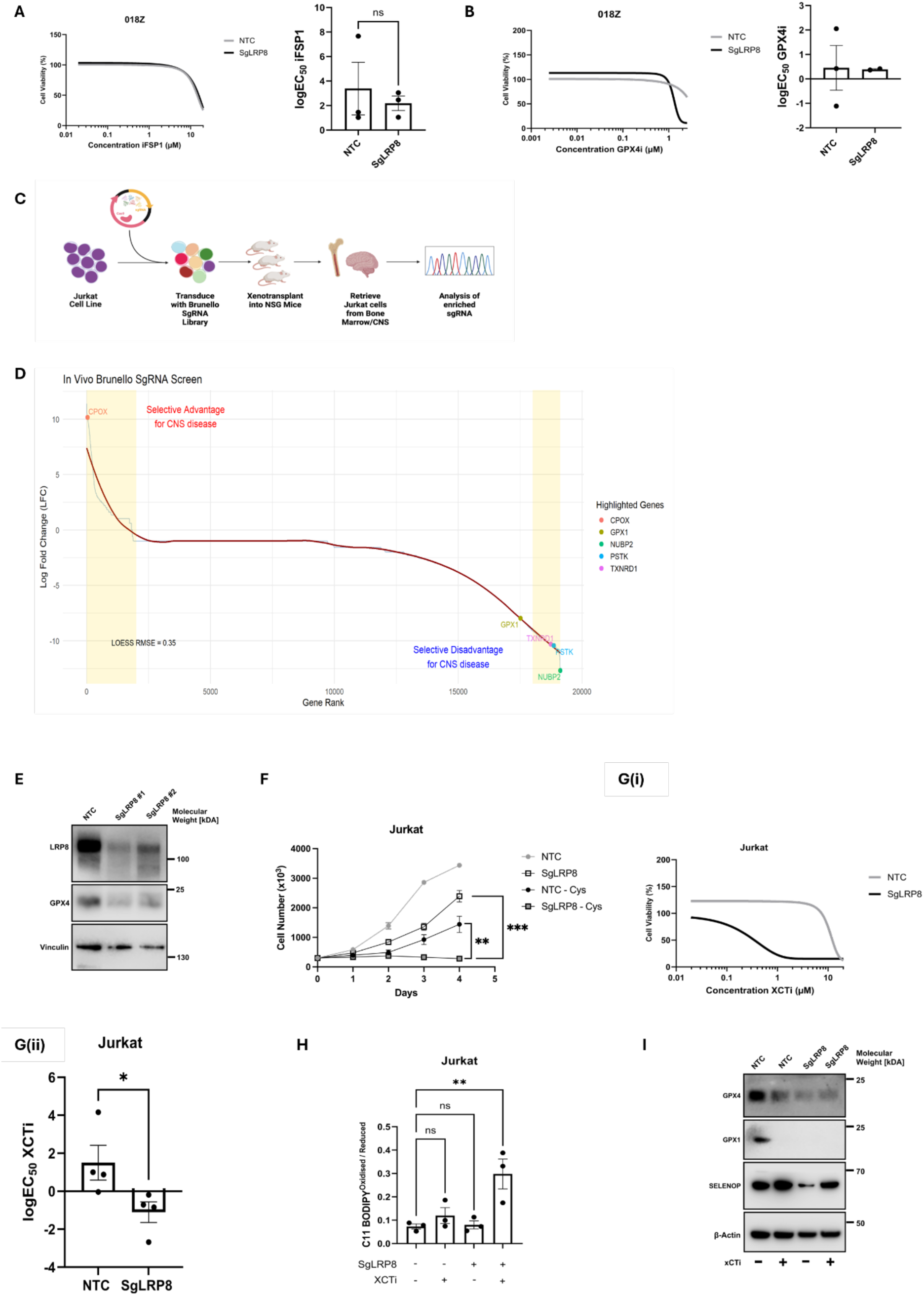
CSF-like Cystine Limiting Conditions are Synthetically Lethal with Selenocysteine Biosynthesis Interference. (**A**) Left Panel: Dose-response curves of 018Z NTC or sgLRP8 #1 cells cultured in RPMI with 10% FBS and treated with iFSP1. Right Panel: logEC_50_ values for iFSP1. Each point indicates the calculated logEC_50_ from independent experiments. P-values calculated using unpaired two-tailed Student’s t-test. Error bars represent mean ± SEM. **(B)** Left Panel: Dose-response curves of 018Z NTC or sgLRP8 #1 cells cultured in RPMI with 10% FBS and treated with RSL3. Right Panel: logEC_50_ values for RSL3. Each point indicates the calculated logEC_50_ from independent experiments (1 sgLRP8 curve could not be fitted due to lack of cytotoxicity). Error bars represent mean ± SEM. **(C)** Schematic representation of the workflow used to screen gene essentiality in the CNS and in the BM with the Brunello CRISPR Cas9 library. Created with BioRender.com. **(D)** Scatterplot showing the rank of the *iron and selenium-*related genes based on the differential score. The regions spanning the most essential and disadvantageous genes for ALL cells in the CNS niche are highlighted in yellow. **(E)** Immunoblot images of LRP8, GPX4 and Vinculin in Jurkat cells transduced with NTC, sgLRP8 1 or sgLRP8 2. **(F)** Cell number of NTC and sgLRP8 #1 cells cultured in media with or without 100 µM L-cystine (Cys) over 96 hours. P-values determined by one-way ANOVA with Dunnett’s multiple comparisons test. N = 3 independent experiments. Error bars represent mean ± SEM. **(G) (i)** Dose-response curves of Jurkat NTC or sgLRP8 #1 cells treated with Erastin in RPMI with 10% FBS. Luminescence values were normalised to vehicle-treated controls. N = 4 independent experiments. **(ii)**: logEC_50_ values for Erastin each data point indicates the calculated logEC_50_ from independent experiments. P-values calculated using unpaired two-tailed Student’s t-test. Error bars represent mean ± SEM. Error bars represent mean ± SEM. **(H)** Ratio of oxidised (510 nm) to reduced (591 nm) BODIPY 581/591 C11 in NTC or sgLRP8 #1 Jurkat cells incubated for 24 hours with or without 1.5 µM Erastin. N = 3 independent experiments. P-values determined by one-way ANOVA with Dunnett’s multiple comparisons test relative to NTC vehicle control. Error bars represent mean ± SEM. **(I)** Immunoblot images of GPX4, GPX1, SELENOP, and β-Actin protein levels in Jurkat sgLRP8 #1 or NTC cells cultured in media with or without 1.5 µM Erastin for 48 hours. ** ≤ 0.01 *** ≤ 0.001

**Supplemental Figure 5:**
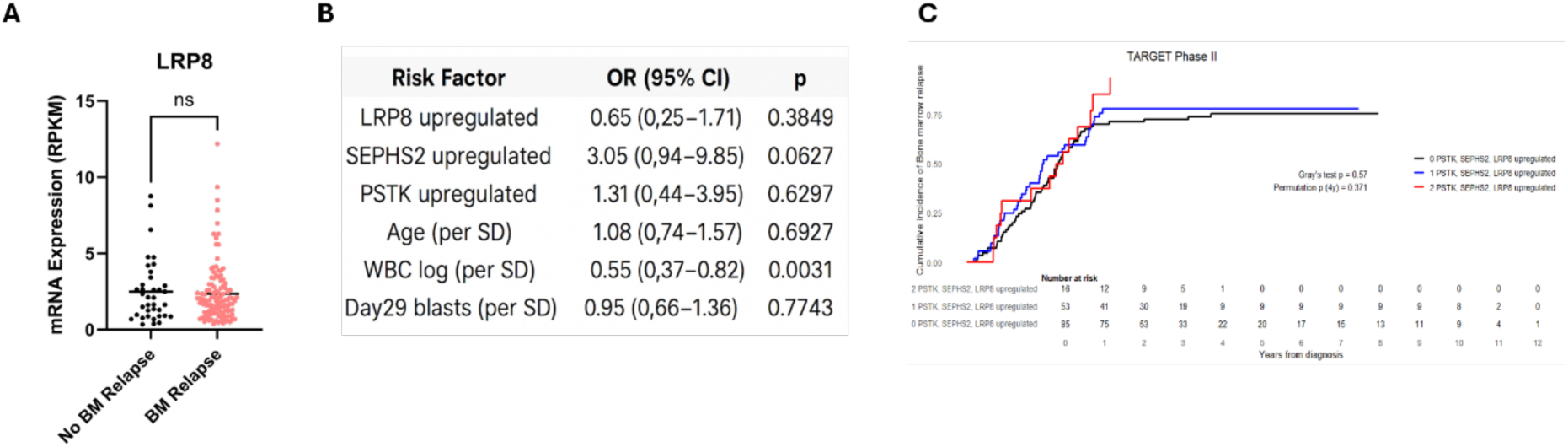
Selenocysteine Biosynthesis Genes Are Not Predictive of Bone Marrow Relapse. **(A)** Diagnostic BM mRNA levels from TARGET Phase II patients (n=203) who relapsed in the BM compared to no BM relapse. P-values calculated using unpaired two-tailed Student’s t-test. Error bars represent mean ± SEM. **(B)** Cox Proportional Hazards model of risk of BM relapse in this dataset with traditional risk factors (MRD positivity of >0.01% at day 29 of induction therapy, WCC > 50 × 10^9^/L, Age >10 years) and upregulation of LRP8, SEPHS2 or PSTK (Z-Score **≥** 1). **(C)** Fine–Gray analysis showing cumulative incidence of bone marrow-involving relapse in patients with upregulation of any one or two or more of LRP8, SEPHS2, or PSTK (gene expression z-score ≥ 1.0).

**Supplemental Figure 6:**
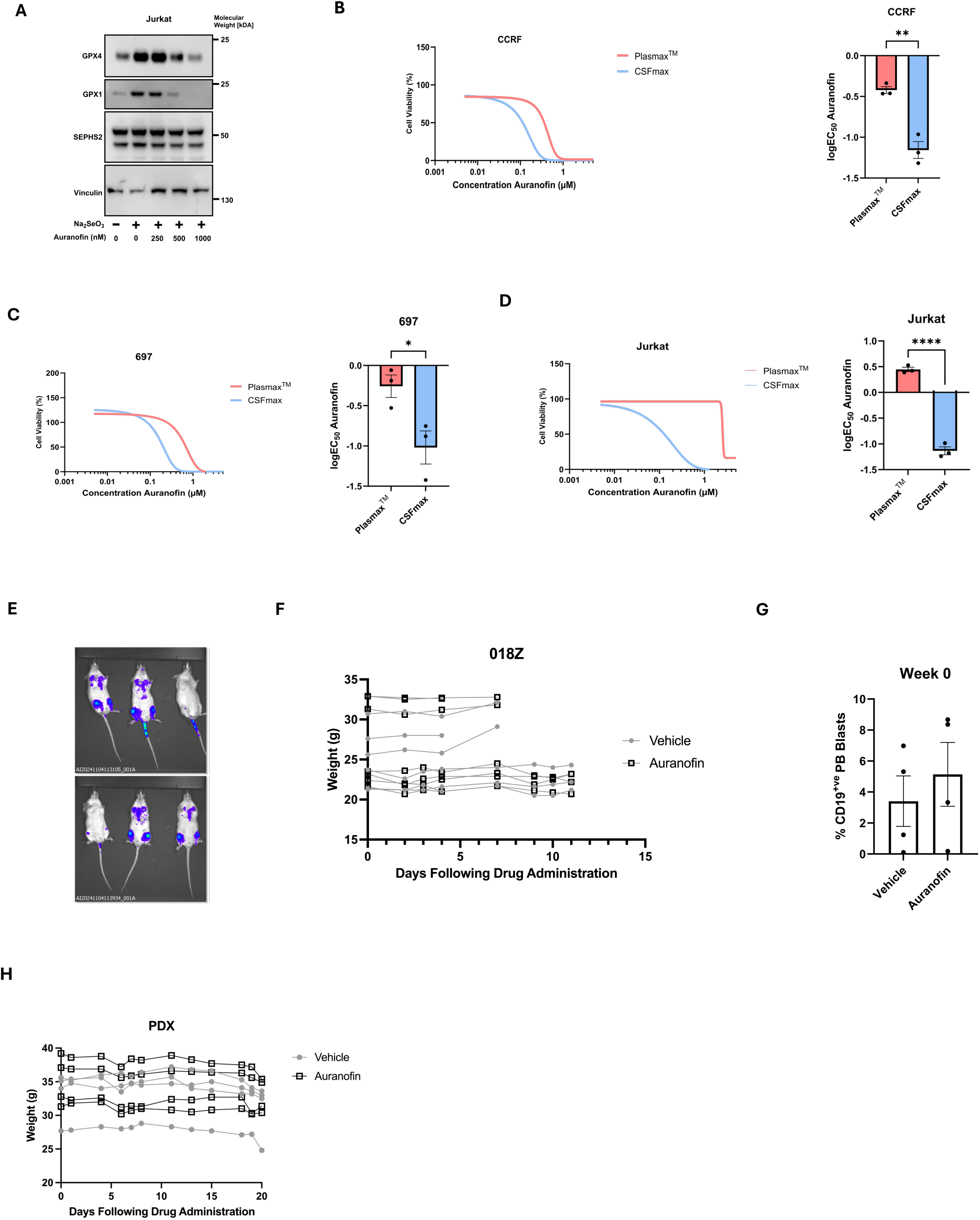
Auranofin as a Pharmacological Strategy to Target the Selenoproteome. (A) Immunoblot image of GPX4, GPX1, SEPHS2 and Vinculin in Jurkat cells treated with 150nM Na_2_SeO_3_ and with an increasing concentration of Auranofin for 72 hours as indicated. The results from 1 experiment representative of 2 are shown. (B-D) Left Panel: Dose-response curves of (B) CCRF, (C) 697, (D) Jurkat cells treated with Auranofin in CSFmax or Plasmax^TM^ as indicated. Luminescence values were normalised to vehicle-treated controls. Points indicate the mean of N = 3 independent experiments. Right Panel: logEC_50_ values for Auranofin. Each point indicates the calculated logEC_50_ from independent experiments (n=3). P-values calculated using unpaired two-tailed Student’s t-test. Error bars represent mean ± SEM (E) Representative images of mice imaged with IVIS 4 days after tail vein injection of 1× 106 luciferase expressing 018Z cells before being assigned to the vehicle (PBS) or treatment (10mg/kg of Auranofin) groups. (F) Mouse weights in vehicle or Auranofin treated cohorts, each line represents an individual mouse. (G) Presence of blasts in the peripheral blood of mice with PDX at treatment start. Error bars represent mean ± SEM. (H) Mouse weights in vehicle and Auranofin treated cohorts, each line represents an individual mouse.

## Notes

### Competing Interest Statement

SaT is the inventor of PlasmaxTM cell culture medium. CH, SaT, EK and VH are named inventors on a patent application for CSFmax cell culture medium (filing date 09/05/2025, patent application number 2507177). All other authors declare no competing financial interests or personal relationships that could have appeared to influence the work reported in this study.

## References

1 Segal, M. B. Extracellular and cerebrospinal fluids. J Inherit Metab Dis 16, 617–638 (1993). 10.1007/bf00711896

2 Iliff, J. J. et al. A paravascular pathway facilitates CSF flow through the brain parenchyma and the clearance of interstitial solutes, including amyloid β. Sci Transl Med 4, 147ra111 (2012). 10.1126/scitranslmed.3003748

3 Spector, R., Robert Snodgrass, S. & Johanson, C. E. A balanced view of the cerebrospinal fluid composition and functions: Focus on adult humans. Experimental Neurology 273, 57–68 (2015). 10.1016/j.expneurol.2015.07.027

4 Halsey, C. & Escherich, G. A “Goldilocks” approach to CNS leukemia is needed. Blood 138, 288–289 (2021). 10.1182/blood.2021011461

5 Schmiegelow, K. et al. Consensus definitions of 14 severe acute toxic effects for childhood lymphoblastic leukaemia treatment: a Delphi consensus. Lancet Oncol 17, e231–e239 (2016). 10.1016/s1470-2045(16)30035-3

6 Gupta, S. et al. Sex-based disparities in outcome in pediatric acute lymphoblastic leukemia: a Children’s Oncology Group report. Cancer 128, 1863–1870 (2022). 10.1002/cncr.34150

7 Savino, A. M. et al. Metabolic adaptation of acute lymphoblastic leukemia to the central nervous system microenvironment is dependent on Stearoyl CoA desaturase. Nat Cancer 1, 998–1009 (2020). 10.1038/s43018-020-00115-2

8 Ferraro, G. B. et al. FATTY ACID SYNTHESIS IS REQUIRED FOR BREAST CANCER BRAIN METASTASIS. Nat Cancer 2, 414–428 (2021). 10.1038/s43018-021-00183-y

9 Ubellacker, J. M. et al. Lymph protects metastasizing melanoma cells from ferroptosis. Nature 585, 113–118 (2020). 10.1038/s41586-020-2623-z

10 Kalkavan, H. et al. Sublethal cytochrome c release generates drug-tolerant persister cells. Cell 185, 3356–3374.e3322 (2022). 10.1016/j.cell.2022.07.025

11 Hassannia, B., Vandenabeele, P. & Vanden Berghe, T. Targeting Ferroptosis to Iron Out Cancer. Cancer Cell 35, 830–849 (2019). 10.1016/j.ccell.2019.04.002

12 Lei, G., Zhuang, L. & Gan, B. The roles of ferroptosis in cancer: Tumor suppression, tumor microenvironment, and therapeutic interventions. Cancer Cell 42, 513–534 (2024). 10.1016/j.ccell.2024.03.011

13 Tobias, A. et al. Breast cancer secretes anti-ferroptotic MUFAs and depends on selenoprotein synthesis for metastasis. bioRxiv, 2023.2006.2013.544588 (2024). 10.1101/2023.06.13.544588

14 Tesfay, L. et al. Stearoyl-CoA Desaturase 1 Protects Ovarian Cancer Cells from Ferroptotic Cell Death. Cancer Res 79, 5355–5366 (2019). 10.1158/0008-5472.Can-19-0369

15 van der Velden, V. H. et al. New cellular markers at diagnosis are associated with isolated central nervous system relapse in paediatric B-cell precursor acute lymphoblastic leukaemia. Br J Haematol 172, 769–781 (2016). 10.1111/bjh.13887

16 Ito, J. et al. PRDX6 dictates ferroptosis sensitivity by directing cellular selenium utilization. Molecular Cell 84, 4629–4644.e4629 (2024). 10.1016/j.molcel.2024.10.028

17 Ellman, G. L. Tissue sulfhydryl groups. Arch Biochem Biophys 82, 70–77 (1959). 10.1016/0003-9861(59)90090-6

18 Persichilli, S., Gervasoni, J., Castagnola, M., Zuppi, C. & Zappacosta, B. A Reversed-Phase HPLC Fluorimetric Method for Simultaneous Determination of Homocysteine-Related Thiols in Different Body Fluids. Laboratory Medicine 42, 657–662 (2011). 10.1309/lmoiah19rg5bkbiq

19 Lu, S. C. Glutathione synthesis. Biochimica et Biophysica Acta (BBA) – General Subjects 1830, 3143–3153 (2013). 10.1016/j.bbagen.2012.09.008

20 Wishart, D. S. et al. HMDB 5.0: the Human Metabolome Database for 2022. Nucleic Acids Res 50, D622–d631 (2022). 10.1093/nar/gkab1062

21 Dixon, Scott J. et al. Ferroptosis: An Iron-Dependent Form of Nonapoptotic Cell Death. Cell 149, 1060–1072 (2012). 10.1016/j.cell.2012.03.042

22 Drummen, G. P., van Liebergen, L. C., Op den Kamp, J. A. & Post, J. A C11-BODIPY(581/591), an oxidation-sensitive fluorescent lipid peroxidation probe: (micro)spectroscopic characterization and validation of methodology. Free Radic Biol Med 33, 473–490 (2002). 10.1016/s0891-5849(02)00848-1

23 Crawford, R. R. et al. Human CHAC1 Protein Degrades Glutathione, and mRNA Induction Is Regulated by the Transcription Factors ATF4 and ATF3 and a Bipartite ATF/CRE Regulatory Element. J Biol Chem 290, 15878–15891 (2015). 10.1074/jbc.M114.635144

24 Dixon, S. J. et al. Pharmacological inhibition of cystine-glutamate exchange induces endoplasmic reticulum stress and ferroptosis. Elife 3, e02523 (2014). 10.7554/eLife.02523

25 Alborzinia, H. et al. LRP8-mediated selenocysteine uptake is a targetable vulnerability in MYCN-amplified neuroblastoma. EMBO Molecular Medicine 15, e18014 (2023). 10.15252/emmm.202318014

26 Zhu, J. et al. Transsulfuration Activity Can Support Cell Growth upon Extracellular Cysteine Limitation. Cell Metab 30, 865–876.e865 (2019). 10.1016/j.cmet.2019.09.009

27 Vande Voorde, J., et al. Improving the metabolic fidelity of cancer models with a physiological cell culture medium. Sci Adv 5, eaau7314 (2019). 10.1126/sciadv.aau7314

28 Kang, J., Ostergaard, J., Wang, X. & Gordon, P. M. Cerebrospinal fluid attenuates the efficacy of methotrexate against acute lymphoblastic leukemia cells. Blood Neoplasia 2, 100057 (2025). 10.1016/j.bneo.2024.100057

29 Alsadeq, A. et al. The role of ZAP70 kinase in acute lymphoblastic leukemia infiltration into the central nervous system. Haematologica 102, 346–355 (2017). 10.3324/haematol.2016.147744

30 Foley, G. E. et al. Continuous culture of human lymphoblasts from peripheral blood of a child with acute leukemia. Cancer 18, 522–529 (1965). 10.1002/1097-0142(196504)18:4<522::AID-CNCR2820180418>3.0.CO;2-J

31 Eckhoff, E. M., Queudeville, M., Debatin, K.-M. & Meyer, L. H. A Novel B Cell Precursor ALL Cell Line (018Z) with Prominent Neurotropism and Isolated CNS Leukemia in a NOD/SCID/huALL Xenotransplantation Model. Blood 114, 1630 (2009). 10.1182/blood.V114.22.1630.1630

32 Frishman-Levy, L. et al. Central nervous system acute lymphoblastic leukemia: role of natural killer cells. Blood 125, 3420–3431 (2015). 10.1182/blood-2014-08-595108

33 Doll, S. et al. FSP1 is a glutathione-independent ferroptosis suppressor. Nature 575, 693–698 (2019). 10.1038/s41586-019-1707-0

34 Soula, M. et al. Metabolic determinants of cancer cell sensitivity to canonical ferroptosis inducers. Nat Chem Biol 16, 1351–1360 (2020). 10.1038/s41589-020-0613-y

35 Kraft, V. A. N. et al. GTP Cyclohydrolase 1/Tetrahydrobiopterin Counteract Ferroptosis through Lipid Remodeling. ACS Cent Sci 6, 41–53 (2020). 10.1021/acscentsci.9b01063

36 Sanson, K. R. et al. Optimized libraries for CRISPR-Cas9 genetic screens with multiple modalities. Nature Communications 9, 5416 (2018). 10.1038/s41467-018-07901-8

37 Heath, A. P. et al. The NCI Genomic Data Commons. Nature Genetics 53, 257–262 (2021). 10.1038/s41588-021-00791-5

38 Talbot, S., Nelson, R. & Self, W. Arsenic trioxide and auranofin inhibit selenoprotein synthesis: implications for chemotherapy for acute promyelocytic leukaemia. British journal of pharmacology 154, 940–948 (2008).

39 Sabatier, P. et al. Comprehensive chemical proteomics analyses reveal that the new TRi-1 and TRi-2 compounds are more specific thioredoxin reductase 1 inhibitors than auranofin. Redox Biology 48, 102184 (2021). 10.1016/j.redox.2021.102184

40 Bak, D. W., Gao, J., Wang, C. & Weerapana, E. A Quantitative Chemoproteomic Platform to Monitor Selenocysteine Reactivity within a Complex Proteome. Cell Chemical Biology 25, 1157–1167.e1154 (2018). 10.1016/j.chembiol.2018.05.017

41 Walz, D. T., DiMartino, M. J., Griswold, D. E., Intoccia, A. P. & Flanagan, T. L. Biologic actions and pharmacokinetic studies of auranofin. The American journal of medicine 75, 90–108 (1983).

42 Madeira, J. et al. Novel protective properties of auranofin: inhibition of human astrocyte cytotoxic secretions and direct neuroprotection. Life sciences 92, 1072–1080 (2013).

43 Mostert, V., Lombeck, I. & Abel, J. A novel method for the purification of selenoprotein P from human plasma. Arch Biochem Biophys 357, 326–330 (1998). 10.1006/abbi.1998.0809

44 Himeno, S., Chittum, H. S. & Burk, R. F. Isoforms of Selenoprotein P in Rat Plasma: EVIDENCE FOR A FULL-LENGTH FORM AND ANOTHER FORM THAT TERMINATES AT THE SECOND UGA IN THE OPEN READING FRAME*. Journal of Biological Chemistry 271, 15769–15775 (1996). 10.1074/jbc.271.26.15769

45 Méplan, C. et al. Relative Abundance of Selenoprotein P Isoforms in Human Plasma Depends on Genotype, Se Intake, and Cancer Status. Antioxidants & Redox Signaling 11, 2631–2640 (2009). 10.1089/ars.2009.2533

46 Williams, M. T. S. et al. The ability to cross the blood–cerebrospinal fluid barrier is a generic property of acute lymphoblastic leukemia blasts. Blood 127, 1998–2006 (2016). 10.1182/blood-2015-08-665034

47 Evans, A. E., Gilbert, E. S. & Zandstra, R. The increasing incidence of central nervous system leukemia in children.(Children’s Cancer Study Group A). Cancer 26, 404–409 (1970).

48 Moore, E. W., Thomas, L. B., Shaw, R. K. & Freireich, E. J. The central nervous system in acute leukemia: a postmortem study of 117 consecutive cases, with particular reference to hemorrhages, leukemic infiltrations, and the syndrome of meningeal leukemia. AMA Archives of Internal Medicine 105, 451–468 (1960).

49 Chi, Y. et al. Cancer cells deploy lipocalin-2 to collect limiting iron in leptomeningeal metastasis. Science 369, 276–282 (2020). 10.1126/science.aaz2193

50 Yoo, S. E. et al. Gpx4 ablation in adult mice results in a lethal phenotype accompanied by neuronal loss in brain. Free Radic Biol Med 52, 1820–1827 (2012). 10.1016/j.freeradbiomed.2012.02.043

51 Leiter, O. et al. Lrp8 knockout mice fed a selenium-replete diet display subtle deficits in their spatial learning and memory function. Behav Neurosci 138, 125–141 (2024). 10.1037/bne0000585

52 Burk, R. F. et al. Deletion of apolipoprotein E receptor-2 in mice lowers brain selenium and causes severe neurological dysfunction and death when a low-selenium diet is fed. J Neurosci 27, 6207–6211 (2007). 10.1523/jneurosci.1153-07.2007

53 Fiskus, W. et al. Auranofin Induces Lethal Oxidative and Endoplasmic Reticulum Stress and Exerts Potent Preclinical Activity against Chronic Lymphocytic Leukemia. Cancer Research 74, 2520–2532 (2014). 10.1158/0008-5472.Can-13-2033

54 Chen, L. et al. Direct inhibition of dioxygenases TET1 by the rheumatoid arthritis drug auranofin selectively induces cancer cell death in T-ALL. J Hematol Oncol 16, 113 (2023). 10.1186/s13045-023-01513-6

55 Natarajan, D., Prasad, N. R., Sudharsan, M., Bharathiraja, P. & Lakra, D. S. Auranofin sensitizes breast cancer cells to paclitaxel chemotherapy by disturbing the cellular redox system. Cell Biochem Funct 41, 1305–1318 (2023). 10.1002/cbf.3865

56 Stafford, W. C. et al. Irreversible inhibition of cytosolic thioredoxin reductase 1 as a mechanistic basis for anticancer therapy. Science Translational Medicine 10, eaaf7444 (2018). doi:10.1126/scitranslmed.aaf7444

57 Giannini, E. H., Brewer, E. J., Jr., Kuzmina, N., Shaikov, A. & Wallin, B. Auranofin in the treatment of juvenile rheumatoid arthritis. Results of the USA-USSR double-blind, placebo-controlled trial. The USA Pediatric Rheumatology Collaborative Study Group. The USSR Cooperative Children’s Study Group. Arthritis Rheum 33, 466–476 (1990). 10.1002/art.1780330402

58 Alexander, S. et al. Impact of Propofol Exposure on Neurocognitive Outcomes in Children With High-Risk B ALL: A Children’s Oncology Group Study. Journal of Clinical Oncology 42, 2671–2679 (2024). 10.1200/JCO.23.01989

59 Cerami, E. et al. The cBio cancer genomics portal: an open platform for exploring multidimensional cancer genomics data. Cancer Discov 2, 401–404 (2012). 10.1158/2159-8290.Cd-12-0095

60 Wishart, D. S. et al. The human cerebrospinal fluid metabolome. J Chromatogr B Analyt Technol Biomed Life Sci 871, 164–173 (2008). 10.1016/j.jchromb.2008.05.001

61 Spector, R. & Johanson, C. E. Sustained choroid plexus function in human elderly and Alzheimer’s disease patients. Fluids Barriers CNS 10, 28 (2013). 10.1186/2045-8118-10-28

62 Sambrook, M. A. The relationship between cerebrospinal fluid and plasma electrolytes in patients with meningitis. Journal of the Neurological Sciences 23, 265–273 (1974). 10.1016/0022-510X(74)90230-5

63 Meseguer, I. et al. Cerebrospinal fluid levels of selenium in patients with Alzheimer’s disease. J Neural Transm (Vienna) 106, 309–315 (1999). 10.1007/s007020050160

64 Zecca, L. et al. Nitrite and nitrate levels in cerebrospinal fluid of normal subjects. J Neural Transm (Vienna) 105, 627–633 (1998). 10.1007/s007020050084

65 Williams, M. T. et al. Interleukin-15 enhances cellular proliferation and upregulates CNS homing molecules in pre-B acute lymphoblastic leukemia. *Blood*, The Journal of the American Society of Hematology 123, 3116–3127 (2014).

66 Lee, N., Carlisle, A. E. & Kim, D. Examining xCT-mediated selenium uptake and selenoprotein production capacity in cells. Methods Enzymol 662, 1–24 (2022). 10.1016/bs.mie.2021.10.002

67 Villar, V. H. et al. Hepatic glutamine synthetase controls N5-methylglutamine in homeostasis and cancer. Nature Chemical Biology 19, 292–300 (2023). 10.1038/s41589-022-01154-9

68 Adams, K. J. et al. Skyline for Small Molecules: A Unifying Software Package for Quantitative Metabolomics. J Proteome Res 19, 1447–1458 (2020). 10.1021/acs.jproteome.9b00640

69 Doench, J. G. et al. Optimized sgRNA design to maximize activity and minimize oa-target effects of CRISPR-Cas9. Nat Biotechnol 34, 184–191, doi:10.1038/nbt.3437 (2016).

